# Lysophosphatidylcholine acyltransferase 1 controls the mitochondrial reactive oxygen species generation and survival of the retinal photoreceptor cells

**DOI:** 10.1101/2021.10.06.463439

**Authors:** Katsuyuki Nagata, Daisuke Hishikawa, Hiroshi Sagara, Masamichi Saito, Sumiko Watanabe, Takao Shimizu, Hideo Shindou

## Abstract

Due to the high energy demands and characteristic morphology, retinal photoreceptor cells require a specialized lipid metabolism for survival and function. Accordingly, dysregulation of lipid metabolism leads to the photoreceptor cell death and retinal degeneration. Mice with a frameshift mutation of lysophosphatidylcholine acyltransferase 1 (*Lpcat1*), which produces saturated phosphatidylcholine (PC) composed of two saturated fatty acids, has been reported to cause spontaneous retinal degeneration (*rd*11 mice). In this study, we performed a detailed characterization of LPCAT1 in the retina and found that genetic deletion of *Lpcat1* induces light-independent and photoreceptor-specific apoptosis in mice. Lipidomic analyses of the retina and isolated photoreceptor outer segment (OS) suggested that loss of *Lpcat1* decreases saturated PC production and affects the proper cellular fatty acid flux, presumably by altering saturated fatty acyl-CoA availabilities. Furthermore, we demonstrated that *Lpcat1* deletion increased mitochondrial reactive oxygen species (ROS) levels in photoreceptor cells, but not in other retinal cells without affecting the OS structure and trafficking of OS-localized proteins. These results suggest that the LPCAT1-dependent production of saturated PC is critical for metabolic adaptation during photoreceptor maturation. Our findings highlight the therapeutic potential of saturated fatty acid metabolism in photoreceptor cell degeneration-related retinal diseases.

## INTRODUCTION

In addition to the *de novo* pathway (Kennedy pathway) of phospholipid biosynthesis (1), fatty acyl moieties of membrane phospholipids are turned over dynamically by de-acylation and re-acylation cycles, which is called the Lands’ cycle (2). Appropriate regulation of phospholipid composition by Lands’ cycles is required for various cellular functions, including lipoprotein production and lipid mediator production (3). In the Lands’ cycle, lysophospholipid acyltransferases (LPLATs) involve fatty acid (FA) re-acylation to generate membrane phospholipid diversity. Depending on their substrate (lysophospholipids and acyl-CoAs) selectivity LPLATs produce specific types of membrane phospholipids, such as polyunsaturated FA (PUFA)-containing phospholipids and saturated FA (SFA)-containing phospholipids.

Among LPLATs, lysophosphatidylcholine acyltransferase 1 (LPCAT1) produces saturated phosphatidylcholine (PC) using lysophosphatidylcholine (LPC) and saturated fatty acyl-CoA, such as palmitoyl-CoA (4-6). Previous studies have shown that the LPCAT1-mediated production of saturated PC is required for the proper functioning of pulmonary surfactant in the lung (6,7) and trafficking of growth factor receptors in cancer cells (8). In addition, *Lpcat1* is essential for the survival of retinal photoreceptor cells (9,10). The critical role of *Lpcat1* in photoreceptor cells was originally uncovered by the analysis of retinal degeneration 11 (*rd*11) mice, possessing a frameshift mutation in the *Lpcat1* gene. In *rd*11 mice, the retinal outer nuclear layer (ONL) composed of photoreceptor cells is rapidly diminished, causing vision loss until 1 month of age.

Photoreceptor cells possess a unique membrane phospholipid composition (11). In the membrane of photoreceptor cells, especially in the outer segment (OS) discs, docosahexaenoic acid (DHA)-containing phospholipids are extremely enriched. Our recent study revealed that the loss of DHA-containing phospholipids leads to the collapse of the OS disc structures and retinal degeneration (12). In parallel with this, significant levels of SFA-containing phospholipids were also present in photoreceptor cells. Therefore, the photoreceptor degeneration in *rd*11 suggests that the LPCAT1-mediated production of SFA-containing phospholipids contributes to photoreceptor survival. However, the molecular basis underlying retinal degeneration in *rd*11 mice remain unclear.

In this study, we investigated the role of LPCAT1 in the retina using the *Lpcat1* knockout (KO) mice. Consistent with *rd*11 mice, the retina of *Lpcat1* KO mice showed a rapid loss of ONL with decreased levels of dipalmitoyl PC (DPPC), the major saturated PC species in the retina. We demonstrated that retinal degeneration in *Lpcat1* KO mice was caused by apoptosis of photoreceptor cells, which occurs in a light-independent manner. Furthermore, *Lpcat1* KO photoreceptor cells accumulated mitochondrial reactive oxygen species (ROS). Our study demonstrated that the roles of LPCAT1 are not only in DPPC production for maintenance of membrane integrity, but also in the regulation of normal mitochondrial functions.

## Results

### Photoreceptor cell apoptosis in *Lpcat1* KO mice

A previous study has shown that a frameshift mutation in the *Lpcat1* gene in *rd*11 and B6-JR2845 mice leads to the spontaneous development of severe retinal degeneration (9). Therefore, to investigate whether the loss of function of LPCAT1 causes retinal degeneration, we used mice genetically deleted for *the Lpcat1* gene. Consistent with *rd*11 and B6-JR2845 mice, a dramatic reduction in ONL and OS thickness was observed in 6-week-old *Lpcat1* KO mice compared to that in *Lpcat1* wild-type (WT) and heterozygous (HZ) mice (Figure 1A and 1B). Meanwhile, the thickness of the inner nuclear layer (INL) and ganglion cell layer (GCL) were similar among the three groups (Figure 1A and 1B). The histological differences between *Lpcat1* WT and HZ mouse retinas were not observed.

**Figure 1.**
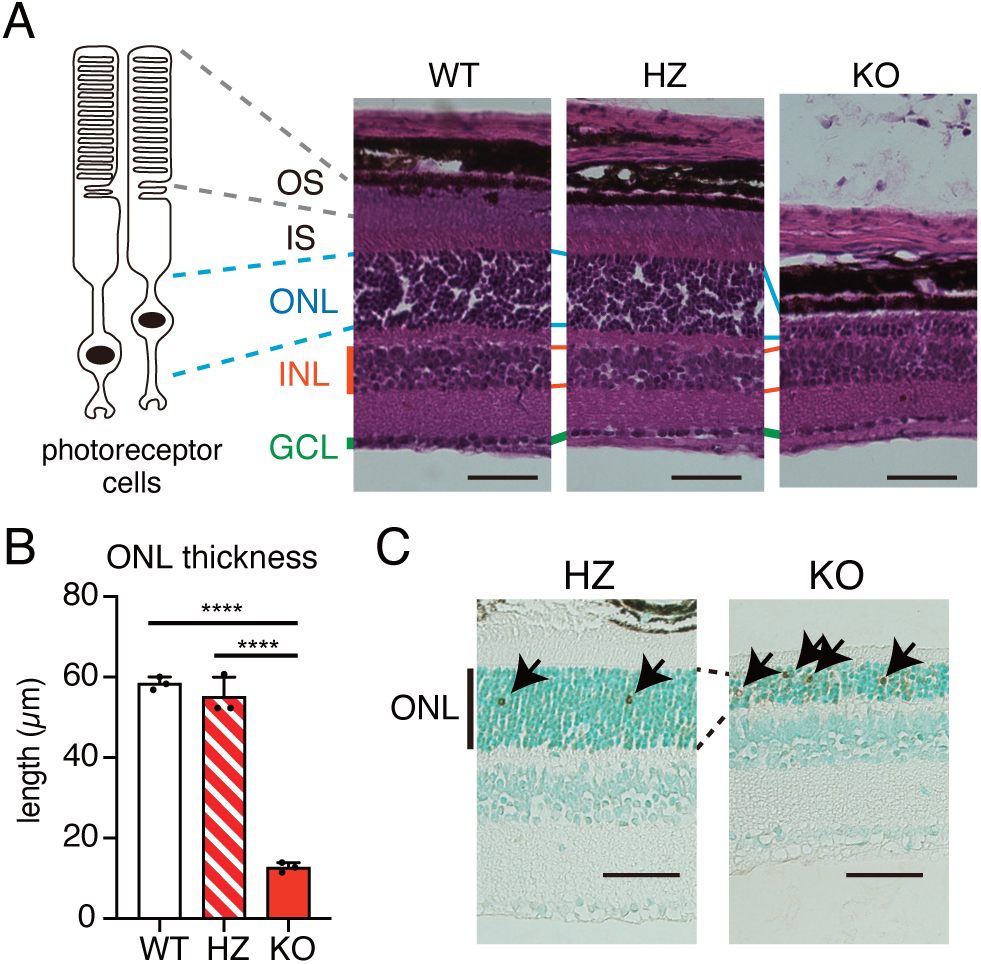
Severe retinal degeneration in *Lpcat1* knockout (KO) mice. A, Histology of *Lpcat1* wild-type (WT), heterozygote (HZ), and KO mouse retina at 6-week-old. Retinal tissue sections were stained with hematoxylin and eosin. The left images illustrate the structure of photoreceptor cells. B, The bar graph showed the outer nuclear layer (ONL) thickness (n=3, for each group). Significance is based on one-way ANOVA followed by Bonferroni’s multiple comparisons test (\*\*\*\**P* < 0.0001). Data are shown as mean + SD of independent experiments. Scale bars are 50 *μ*m. C, Representative images of terminal deoxynucleotidyl transferase dUTP nick-end labeling (TUNEL) staining of 4-week-old *Lpcat1* HZ and KO retina (n=3, for each group). Nuclei were stained with Methyl Green. Arrows indicated TUNEL-positive nuclei detected by 3,3’-diaminobenzidine (DAB, shown in brown). Scale bars are 50 *μ*m. INL, inner nuclear layer; GCL, ganglion cell layer; OS, outer segment; and IS, inner segment.

Apoptosis of photoreceptor cells is a major cause of retinal degeneration (13). To assess the cause of retinal degeneration in *Lpcat1* KO mice, we visualized the apoptotic cells in the retina using terminal deoxynucleotidyl transferase dUTP nick-end labeling (TUNEL). TUNEL analyses indicated that the number of apoptotic photoreceptor cells was higher in 4-week-old *Lpcat1* KO mice than in HZ mice (Figure 1C). Notably, the increase in apoptotic cells in global *Lpcat1* KO mice was not observed in the lungs, liver, or other retinal cell layers (Figure 1C and Figure S1A and S1B). These results indicate that loss of *Lpcat1* leads to photoreceptor cell apoptosis in a specific manner.

### Onset of the photoreceptor apoptosis in *Lpcat1* KO mice

To assess the onset of photoreceptor cell apoptosis in *Lpcat1* KO mice, we performed TUNEL staining of the retina at different time points. Apoptotic cells in the retina of *Lpcat1* KO mice increased greatly from postnatal day 8 (P8) (Figure 2A). Consistent with this, the thickness of ONL was thinner in *Lpcat1* KO mice at P8, while no difference in apoptotic cell number and ONL thickness was seen in the P6 retina (Figure 2A and 2B). Dramatic increases in the expression of *rhodopsin*, critical for phototransduction and OS formation (14), at this time point (Figure 2C), suggests that photoreceptor apoptosis in *Lpcat1* KO retinas occurred concomitantly with photoreceptor OS formation.

**Figure 2.**
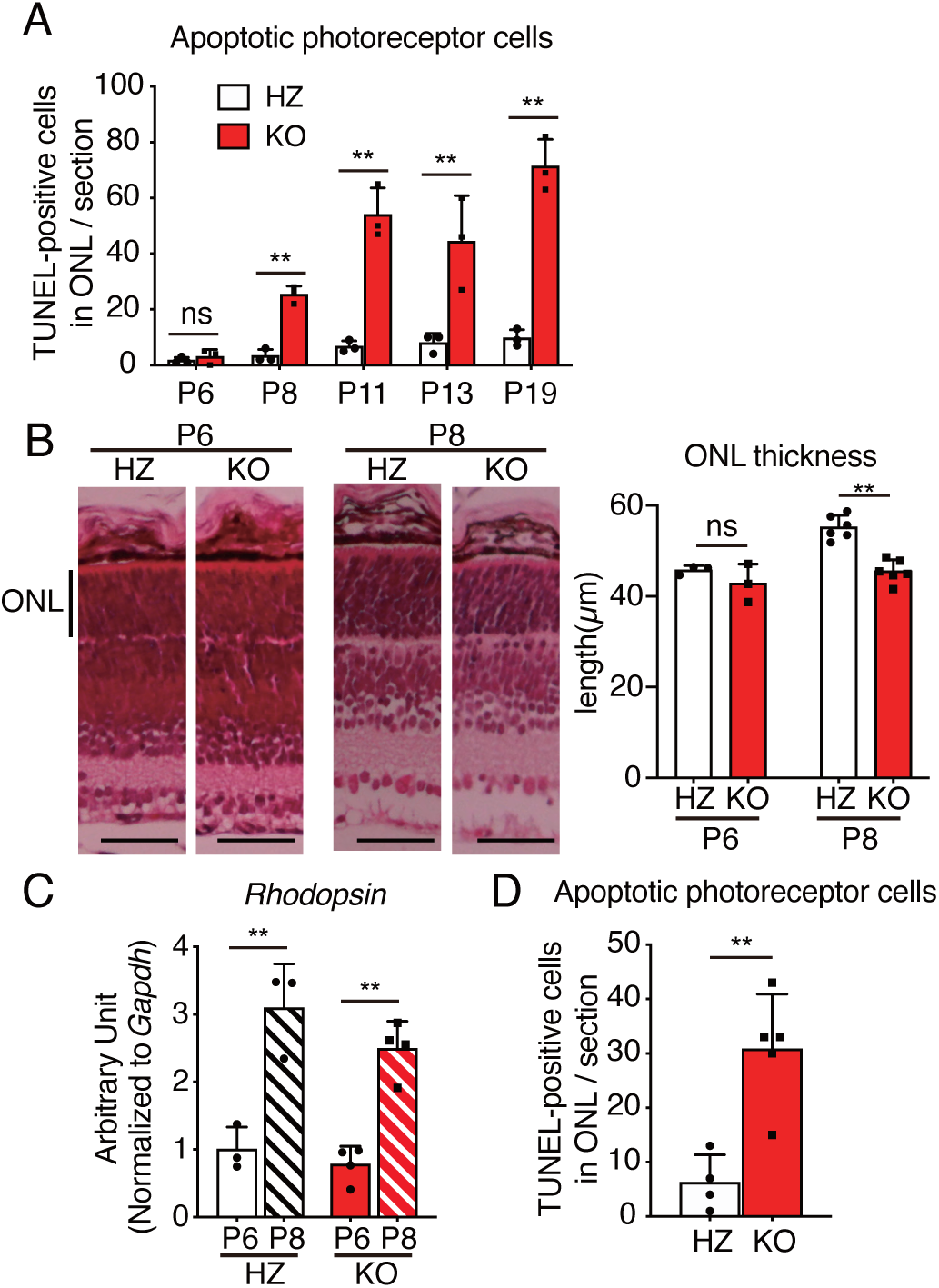
Onset of photoreceptor apoptosis in *Lpcat1* knockout (KO) mice. A, Total numbers of apoptotic cells in the retinal section of postnatal day 6 (P6), P8, P11, P13 and P19 *Lpcat1* heterozygote (HZ) and KO outer nuclear layer (ONL) (n=3). B, The ONL thickness in P6 and P8 *Lpcat1* HZ and KO mice. The left images showed the retinal structure stained with hematoxylin and eosin. The right bar graph showed the ONL thickness (P6, n=3 for each group; P8, n=6 for each group). Scale bars are 50 *μ*m. C, The levels of Rhodopsin mRNA in P6 and P8 of *Lpcat1* HZ and KO retina. The expression level was normalized by glyceraldehyde 3-phosphate dehydrogenase (*Gapdh*) (n=3 for HZ, n=4 for KO). D, Numbers of apoptotic photoreceptor cells in P9 dark-reared *Lpcat1* HZ and KO retina (n=4 for HZ, n=5 for KO). A-C, significance is based on two-way ANOVA followed by Bonferroni’s multiple comparisons test (\*\**P* < 0.01, ns; no significance). Data are shown as mean + SD of independent experiments. D, Significance is based on unpaired *t*-test (\*\**P* < 0.01). Data are shown as mean + SD of independent experiments.

As OS is required for phototransduction, we hypothesized that photoreceptor cell death in *Lpcat1* KO mice is related to light exposure. Thus, we investigated the apoptosis of photoreceptor cells in P9 mice, which raised in complete darkness from P2 to P9. However, photoreceptor cell apoptosis was observed even in dark-reared *Lpcat1* KO mice (Figure 2D), suggesting that retinal degeneration in *Lpcat1* KO mice was independent of light stimulation.

### Altered phospholipid composition in *Lpcat1* KO mouse retina

Photoreceptor cells undergo dynamic morphological and metabolic alterations during retinal maturation (15). Since LPCAT1 produces SFA-containing PC species, we next analyzed PC composition and LPCAT1 expression during retinal maturation. As shown in Figure 3, liquid chromatography-tandem mass spectrometry (LC-MS/MS) analysis showed a gradual alteration in retinal PC composition during maturation. Consistent with previous studies (11,12), the proportion of DHA-containing PC species, including PC 38:6, PC 40:6, and PC 44:12, increased along with retinal maturation (Figure 3). Coinciding with the onset of photoreceptor cell apoptosis in *Lpcat1* KO, DPPC, a major product of LPCAT1, was increased in the retina of WT mice between 1 and 2 weeks of age (Figure 3). Consistent with the substrate selectivity of LPCAT1 *in vitro* (4,6), DPPC levels were decreased by half in *Lpcat1* KO retinas compared to WT retinas. Thus, DPPC production in the retina largely depends on LPCAT1 expression. However, because the induction of LPCAT1 mRNA and protein was not observed during retinal maturation (Figure S2A and S2B), post-translational modifications or the increased substrate supply for LPCAT1, such as LPC and palmitoyl-CoA, may affect the age-dependent elevation of DPPC.

**Figure 3.**
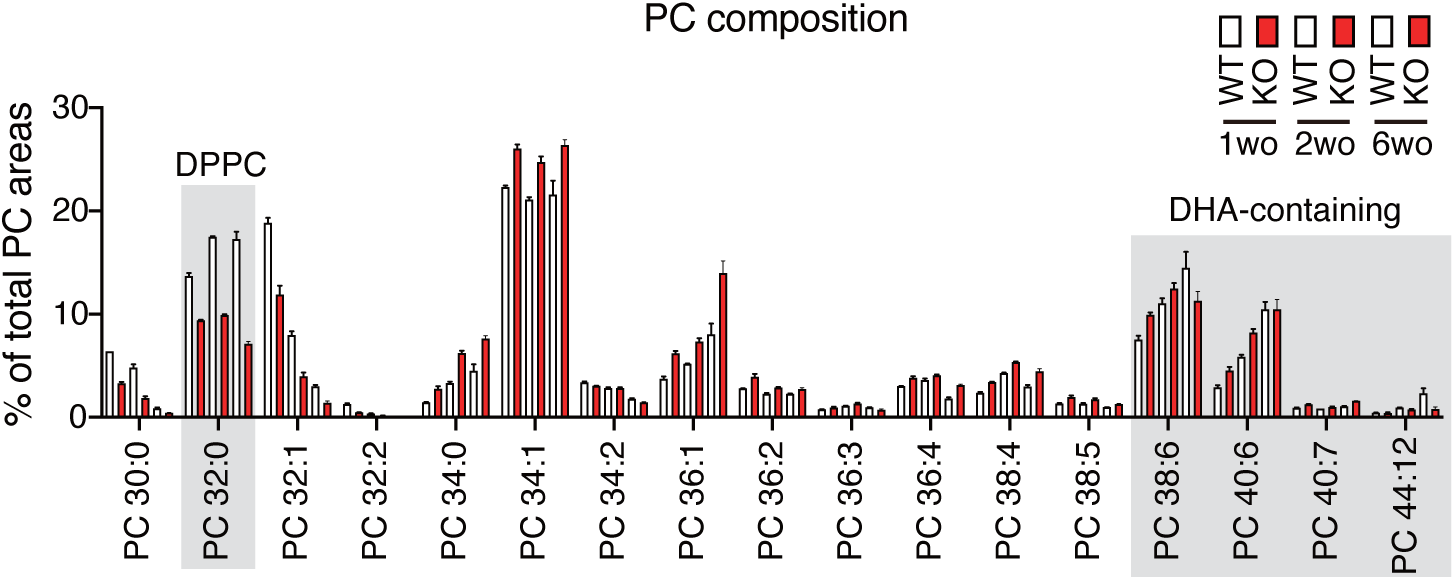
Loss of LPCAT1 leads to the alteration of the retinal phosphatidylcholine (PC) composition. PC composition of the retina. Retinas were prepared from 1-, 2- and 6-week-old (wo) *Lpcat1* wild-type (WT) and knockout (KO) mice (n=3 for each group). Data are shown as mean + SD of independent experiments.

### Transcriptomic analysis of *Lpcat1* KO retina and isolated photoreceptor cells

Next, we aimed to clarify the cause of photoreceptor cell death in *Lpcat1* KO mice by comparing the mRNA expression profiles with the control mice. First, we analyzed the transcriptomic differences in the 2-week-old retinas of *Lpcat1* WT and KO mice (Figure 4A and 4B). Gene groups that increased significantly in *Lpcat1* KO retinas were analyzed by functional classification using “the Database for Annotation, Visualization, and Integrated Discovery (DAVID).” These results suggest that many of the genes upregulated in *Lpcat1* KO retinas were related to immune and inflammatory responses (Figure 4B and Table S1). In addition, we found that the expression of genes termed immediate early genes (IEGs), such as *Fosb, Fos, Egr1*, and *Egr2*, was higher in *Lpcat1* KO than in the control retina (Table S1). Among them, we found that *Fosb* expression in *Lpcat1* KO was significantly higher than in the control retinas, even in dark-reared mice (Figure 4C and 4D). As IEGs are rapidly upregulated by various cellular stimuli, particularly in neurons (16), we performed qPCR analyses using isolated a photoreceptor cell marker CD73-positive and -negative cells (17) from the retina to identify the cellular sources of their upregulation. The qPCR results showed increased expression of IEGs in CD73 negative cells (Figure 4E and Figure S3A). The induction of *Fosb* in *Lpcat1* KO retinas was slightly delayed from the onset of retinal degeneration (Figure 4C), suggesting that it might be triggered by apoptosis of photoreceptor cells.

**Figure 4.**
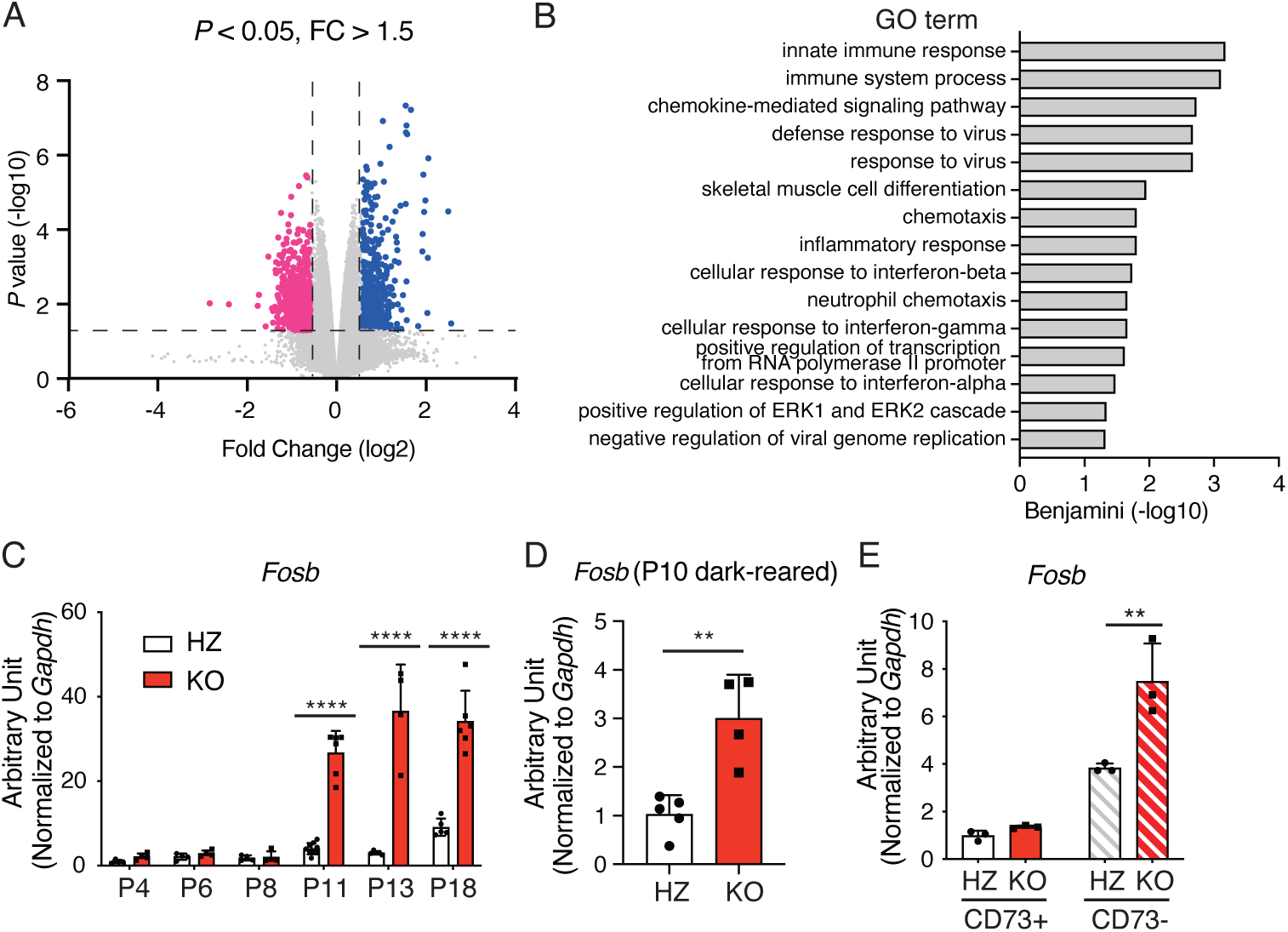
Transcriptomic analysis of *Lpcat1* knockout (KO) retina. A, Volcano plot of differentially expressed genes (DEGs) between postnatal day 14 (P14) *Lpcat1* wild-type (WT) and KO retina. DEGs (Fold change > 1.5 or < -1.5, *P* < 0.05) are highlighted in blue; increased in *Lpcat1* KO, and magenta; decreased in *Lpcat1* KO (n=4 for each group). B, Functional annotation of upregulated genes (shown in blue in Figure 4A) in *Lpcat1* KO retina. C-E, The expression of *Fosb* during neonatal development (C, n=4 for HZ and n=9 for KO), in P10 dark-reared (D, n=4 for HZ and n=5 for KO), and in CD73-positive (photoreceptor cells) or -negative cells (E, n=3 for each group) of *Lpcat1* heterozygote (HZ) and KO retina. The expression level was normalized by glyceraldehyde 3-phosphate dehydrogenase (*Gapdh*). Data are shown as mean + SD of independent experiments. C and E, significance is based on two-way ANOVA followed by Bonferroni’s multiple comparisons test (\*\**P* < 0.01, \*\*\*\**P* < 0.0001). D, significance is based on unpaired *t*-test (\*\**P* < 0.01).

To clarify the primary cause of photoreceptor cell apoptosis in *Lpcat1* KO mice, we performed microarray analysis using isolated photoreceptor cells at P8 when apoptosis of photoreceptor cells in *Lpcat1* KO mice begins. As a result of functional classification analysis, no significant enrichment of the biological process altered in *Lpcat1* KO photoreceptor cells was found in the gene list, except for the categories of cell adhesion and melanin biosynthetic process (Figure S3B and S3C). Since most of the downregulated genes listed in these categories in *Lpcat1* KO photoreceptor cells were reported to be highly expressed in retinal pigment epithelial (RPE) cells, it seemed to reflect the reduced contamination of RPE cell fragments in isolated photoreceptor cells in *Lpcat1* KO mice. This could be caused by the loss of cell-cell tight interactions triggered by photoreceptor death. Indeed, the qPCR analysis showed that mRNA expression of RPE65, an RPE-specific gene, was lower in isolated photoreceptor cells of *Lpcat1* KO mice than in control mice (Figure S3D).

A previous study reported that the failure of SFA-containing PC production due to loss of LPCAT1 leads to excessive polyunsaturated FA accumulation and endoplasmic reticulum (ER) stress response in the retina (18). However, under the present assay conditions, no induction of CCAAT-enhancer-binding protein homologous protein (*Chop*), an ER stress marker, was observed both in *Lpcat1* KO retina and isolated photoreceptor cells (Figure S3E).

### Trafficking and functions of OS localized proteins in *Lpcat1* KO mice

Based on the microarray analyses showing no obvious causative changes in photoreceptor cell death in *Lpcat1* KO mice, we investigated whether the altered membrane phospholipid composition in *Lpcat1* KO photoreceptor cells influences the localization and functions of proteins essential for photoreceptor survival. Consistent with the previous observation of *Lpcat1* mutant mice (9), the structure of *Lpcat1* KO OS discs photoreceptor cells appeared normal (Figure 5A). We then explored the possibility that an altered PC composition of *Lpcat1* KO photoreceptor OS membrane affected the localization and function of phototransduction-related proteins, since the mislocalization of photoreceptor OS localized proteins, such as rhodopsin and phosphodiesterase 6β (PDE6β), is observed in various types of retinal degeneration (19,20). In parallel with the DHA-containing PCs enriched in the OS discs (12), we found that DPPC was also abundant in the OS and decreased by half in *Lpcat1* KO (Figure 5B). This result suggested that the LPCAT1-mediated production of DPPC contributes to maintaining proper PC composition in the photoreceptor OS membrane.

**Figure 5.**
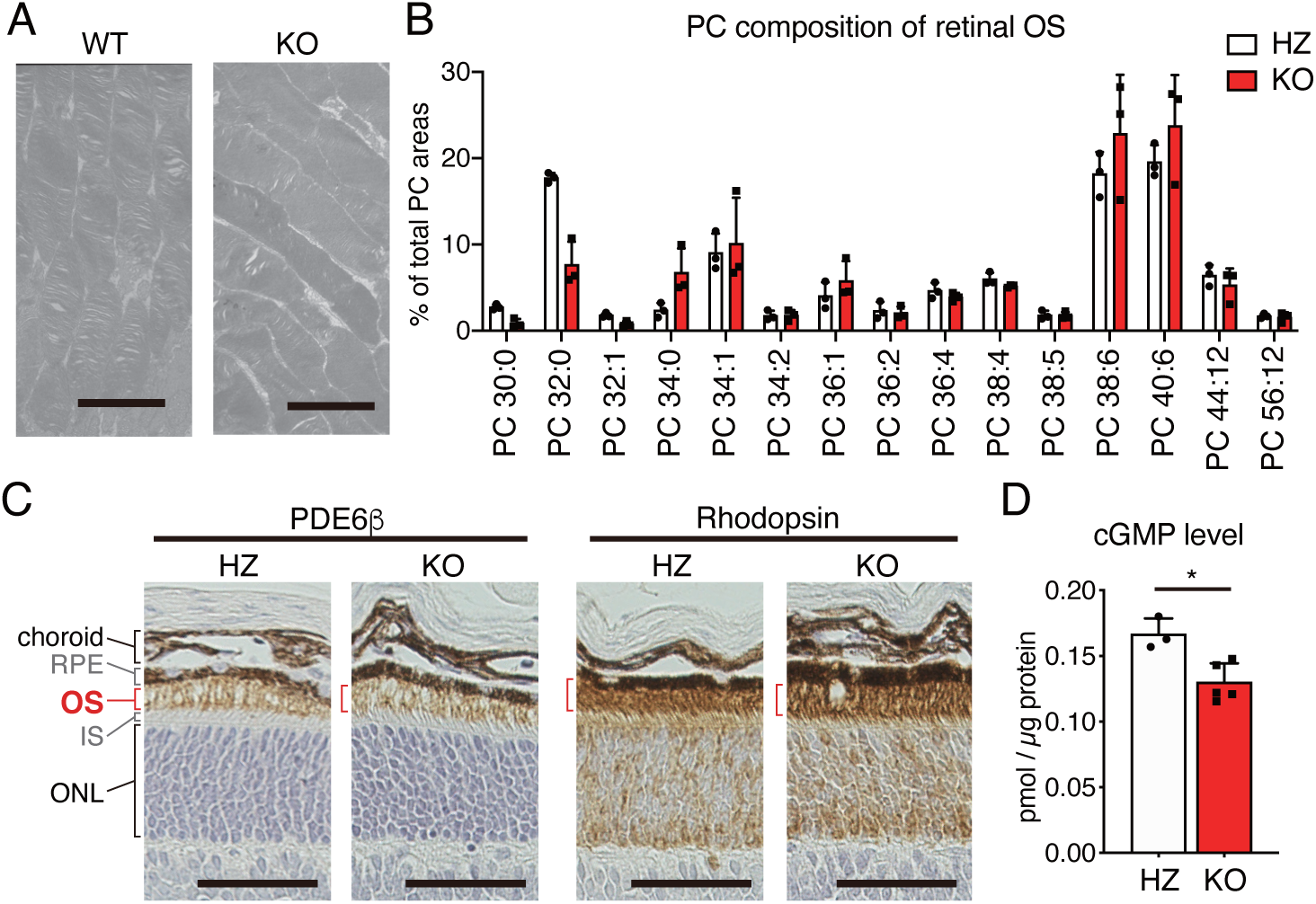
Effect of LPCAT1 deficiency on the structure and function of photoreceptor cells. A, Representative transmission electron microscopy (TEM) images of photoreceptor outer segment (OS) in 4-week-old *Lpcat1* wild-type (WT) and knockout (KO) mice (n=3, for each group). Scale bars are 3 *μ*m. B, Fatty acid composition of phosphatidylcholine (PC) in the isolated photoreceptor OS. OS samples were prepared from postnatal day 19 (P19) *Lpcat1* heterozygote (HZ) and KO mice (n=3 for each group). Data are shown as mean + SD of independent experiments. C, Representative images of PDE6β and Rhodopsin in P13 *Lpcat1* HZ and KO mice by immunohistochemistry analysis. Signals were detected by 3,3’-diaminobenzidine (DAB, shown in brown), observed in the OS region (shown in red brackets). The dark brown colors in RPE and choroid were DAB-independent (n=3 for each group). Scale bars are 50 *μ*m. D, cGMP level in P11 *Lpcat1* HZ and KO retina (n=3 for HZ, n=5 for KO). Data are shown as mean + SD of independent experiments. Significance is based on unpaired *t*-test (\**P* < 0.05). RPE, retinal pigment epithelium; OS, outer segment; IS, inner segment; and ONL, outer nuclear layer.

As we found that *Lpcat1* deletion leads to altered OS membrane PC composition, we next investigated the localization of rhodopsin and PDE6β in the retina. Unexpectedly, both rhodopsin and PDE6β were normally localized in the photoreceptor OS in *Lpcat1* KO mice, despite the altered membrane PC composition in the OS (Figure 5C). PDE6β is a rod photoreceptor-specific subunit of PDE that controls cyclic nucleotide-gated (CNG) channel-mediated cation influx into photoreceptor cells (21). Mice harboring the loss-of-function mutant of PDE6β (termed *rd*1 mouse) showed marked cGMP accumulation and subsequent severe retinal degeneration (22,23). As *rd*1 mice show similar phenotypes to those of *Lpcat1* KO, early-onset and light-independent retinal degeneration (20), we assessed whether the altered membrane lipid composition in *Lpcat1* KO photoreceptor cells leads to PDE6β dysfunction. However, cGMP levels in the retina were slightly decreased rather than increased in *Lpcat1* KO mice compared to control mice (Figure 5D). Since the accumulated cGMP-dependent increase in calcium ion influx triggers retinal degeneration in *rd*1 mice, the underlying mechanisms of retinal degeneration in *Lpcat1* KO mice differ from those in PDE6β mutated mice (*rd*1).

### Increased mitochondrial oxidative stress in *Lpcat1* KO mice

Excess SFA induces apoptosis (24). Similarly, stearoyl-CoA desaturase (SCD), which converts saturated fatty acyl-CoA to monounsaturated fatty acyl-CoA, protects cells from SFA-induced cell death (25). Together with the significant decrease in the PC 32:1/PC 32:0 ratio (Figure S4A) and SCD mRNA (Figure S4B) during photoreceptor maturation, mature photoreceptor cells are presumably less potent in reducing the intracellular SFA levels by the unsaturation. We hypothesized that SFA stress is involved in photoreceptor-specific apoptosis in *Lpcat1* KO mice based on these results. Several factors, including ceramide accumulation, mitochondrial reactive oxygen species (ROS) production, and ER stress, trigger SFA-induced apoptosis (26-29). Since our transcriptomic analyses showed no sign of increased ER stress in *Lpcat1* photoreceptor cells (Figure S3E), we postulated that the altered palmitoyl-CoA flux (Figure 6A) by *Lpcat1* deletion might be related to retinal degeneration.

**Figure 6.**
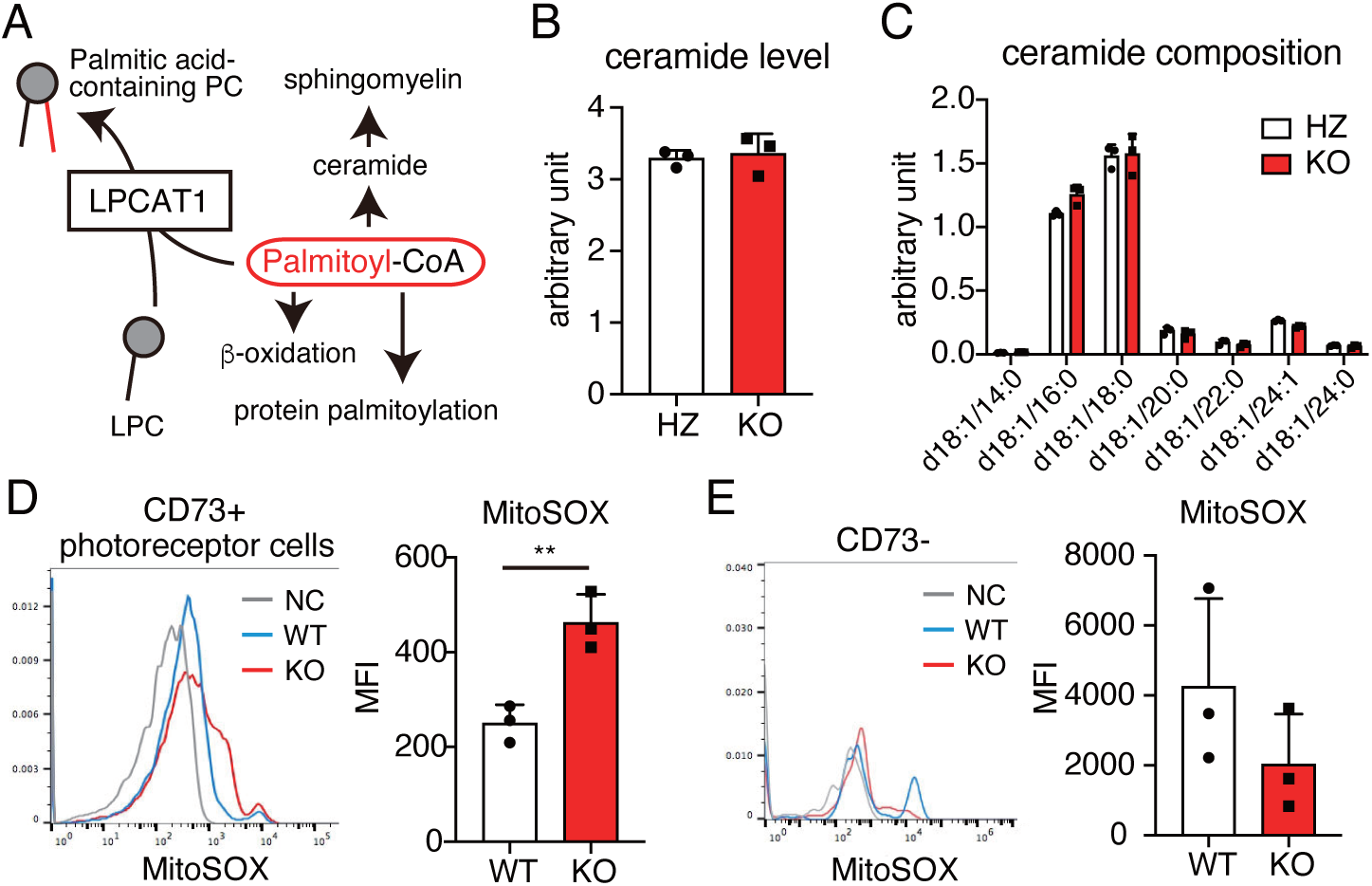
Increased mitochondrial superoxide production in *Lpcat1* knockout (KO) photoreceptor cells. A, Intracellular metabolic pathways of palmitoyl-CoA. B and C, Total area values of ceramides (B) and ceramide composition (C) in P11 *Lpcat1* heterozygote (HZ) and KO retina (n=3 for each group). B, The sum of area values of ceramides were normalized by protein amount and the area values of the internal standard (d18:1/12:0-ceramide) (represented as units/pmol internal standard/*μ*g protein). Data are shown as mean + SD of independent experiments. D and E, Analysis of mitochondrial superoxide production by 3-week-old *Lpcat1* wild-type (WT) and KO photoreceptor cells (D) or retinal cells other than photoreceptor cells (E) (n=3 for each group). Retinal cells were stained with CD73 (photoreceptor cells) and MitoSOX Red (mitochondrial superoxide). Data are shown as mean + SD of independent experiments. Significance is based on unpaired *t*-test (\*\**P* < 0.01). MFI, mean fluorescence intensity.

To this end, we determined whether the *Lpcat1* deletion leads to ceramide accumulation in the retina. A slight increase in sphingomyelin (SM) suggested that loss of *Lpcat1* increased palmitoyl-CoA availability for sphingolipid synthesis. However, we could not find any clear differences in ceramide levels and compositions between genotypes (Figure S4C, Figure 6B and 6C).

Finally, we explored the possibility that the *Lpcat1* deletion leads to increased mitochondrial FA β-oxidation and consequent ROS accumulation in mature photoreceptor cells. We analyzed the mitochondrial ROS accumulation in retinal cells of P21 control and *Lpcat1* KO mice. Flow cytometric analyses of retinal CD73 positive (photoreceptor) cells showed that the ROS accumulated photoreceptor cells in *Lpcat1* KO mice were increased compared to the control mice (Figure 6D and Figure S5A-C). Consistent with TUNEL staining, we observed no differences in ROS accumulation between *Lpcat1* KO and control CD73 negative cells (Figure 6E). These results suggest that mitochondrial ROS accumulation triggered by *Lpcat1* deletion is a photoreceptor-specific phenomenon contributing to apoptosis induction.

## DISCUSSION

Here, we demonstrated that the loss of *Lpcat1* leads to an early onset of severe retinal degeneration, triggered by light stimulus-independent photoreceptor cell apoptosis. Although photoreceptor cell damage induces various types of retinal degeneration, the molecular basis of photoreceptor cell death is not fully understood. Our present study suggests that *Lpcat1* deletion-triggered disruption of palmitoyl-CoA flux leads to mitochondrial ROS accumulation and denaturation in photoreceptor cells.

FA saturation of membrane phospholipids affects membrane fluidity and flexibility through their biophysical properties (30,31). A recent study showed that PUFA-containing phospholipids, especially in DHA-containing phospholipids, are enriched in the center of photoreceptor discs. In contrast, SFA-containing species are enriched in the rim region (32). Therefore, significant levels of saturated PCs in the OS membrane suggest that the photoreceptor OS membrane and/or OS-localized protein require the saturated PC-enriched membrane for their normal structure and functions. In the case of DHA-containing phospholipids, it has been reported that the dramatic decrease in these phospholipids in 1-acylglycerol-3-phosphate *O*-acyltransferase 3 (AGPAT3), also known as lysophosphatidic acid acyltransferase 3 (LPAAT3), KO retina cause the collapse of photoreceptor discs, suggesting the importance of the FA composition of photoreceptor discs in maintaining their structures (12). However, in contrast, the structure of *Lpcat1* KO photoreceptor discs appeared to be normal, regardless of a significant decline in DPPC and PC 30:0. Thus, an increased proportion of PC 34:0, another type of SFA-containing PC, in *Lpcat1* KO photoreceptor OS, may play redundant roles, at least in the formation and/or maintenance of OS disc structures. Conversely, cGMP levels in *Lpcat1* KO retinas were lower than those in WT mice (Figure 5D). Although the decreased cGMP levels in the *Lpcat1* KO retina may reflect the lower number of photoreceptor cells in these mice, it is also possible that the PDE6β activity was affected by altered PC composition in photoreceptor OS membrane. The detailed mechanisms of this observation should be clarified in future studies.

We demonstrated that mitochondrial ROS accumulation was observed in *Lpcat1* KO photoreceptor cells. This suggests that they are highly susceptible to mitochondrial ROS; therefore, mitochondrial FA oxidation should be strictly regulated. A recent study revealed that photoreceptor cells depend largely on fatty acid oxidation in the mitochondria for ATP production (15,33). Together with the fact that mature photoreceptor cells require high levels of ATP to maintain the depolarized state in darkness using ATP-dependent channels (34), LPCAT1-mediated incorporation of saturated fatty acids into PC may be required not only for the production of desaturated PCs but also for the proper control of mitochondrial FA oxidation. Since knockdown of *Lpcat1* expression leads to tumor cell death (8), *Lpcat1*-dependent cell survival of mature photoreceptor cells and cancer cells may show some metabolic similarities.

SCDs are also related to controlling cellular SFA levels by changing saturated acyl-CoA to monounsaturated fatty acyl-CoA (35). It is reported to protect cells from SFA-induced mitochondrial ROS accumulation and apoptosis by changing the fate of SFA from mitochondria to lipid droplets (36,37). Therefore, a dramatic decrease in SCD expression during photoreceptor maturation, which is reflected in the decreased PC 32:1/PC 32:0 (DPPC) ratio, may also be associated with the sensitivity of *Lpcat1* deletion-triggered increase in palmitoyl-CoA availability. It is also possible that altered mitochondrial membrane lipid composition directly affects mitochondrial functions, such as electron transfer, permeability, and fusion/fission. Additionally, we could not exclude the possibility that mitochondrial denaturation and ROS accumulation result from photoreceptor apoptosis. Therefore, further studies are required to decipher the molecular mechanisms of mitochondrial ROS accumulation in *Lpcat1* KO photoreceptor cells.

From the transcriptomic analyses, we identified the apparent induction of FosB mRNA in the *Lpcat1* KO retina during the progression of retinal degeneration. Although the exact roles of FosB in the retina are unclear, it has been reported that *Fosb* is induced in the retina during retinal degeneration (38). As the onset of photoreceptor cell death was followed by the upregulation of *Fosb* in *Lpcat1* KO mice, this might not be the cause of retinal degeneration. However, considering the role of FosB as a component of activator protein-1 (AP-1), a dimeric transcription factor involved in inflammation, angiogenesis, and apoptosis (39,40), our results suggest that *Fosb* inhibition may be involved in retinal degeneration in *Lpcat1* KO mice. Importantly, the upregulation of *Fosb* is also found in the monocytes of patients with age-related macular degeneration (AMD) (41). Therefore, cell types that upregulate *Fosb* during retinal degeneration in *Lpcat1* KO retinas will be clarified by single-cell RNA sequencing in a future study.

In summary, we presented a detailed analysis of *Lpcat1* deletion-induced retinal degeneration. Our data suggest that LPCAT1 is required not only for SFA-containing PC production but also for regulating lipid metabolism by controlling the amount of palmitoyl-CoA in photoreceptor cells. Increasing evidence indicates that disrupted lipid metabolism is the cause of photoreceptor cell death. Therefore, our data provide further insights into the mechanisms underlying photoreceptor cell death-triggered retinal degeneration.

## EXPERIMENTAL PROCEDURES

### Animals

All animal experiments were approved and performed following the guidelines of the Animal Research Committee of the National Center for Global Health and Medicine (12053, 13009, 14045, 15037, 16062, 17054, 18046, 19039, 20037, and 21083).

### Mice

*Lpcat1* KO mice (C57BL6/N background) generated in our previous study were used in this study (6). C57BL6/N mice were reported to show retinal degeneration because they harbor a mutation in *the Crumbs homolog 1* (*Crb1*) gene, termed *rd*8 (23,42). Thus, we crossed *Lpcat1* KO mice with C57BL6/J mice to remove the *rd*8 mutation. Mice without the *rd*8 homozygous mutation were used in this study. Mice were reared in a normal 12 hours light/dark cycle or complete darkness from P2 to P9.

### Histological analysis of mouse retina

For frozen sections, enucleated eyes were fixed in 4% paraformaldehyde for 2 h and washed three times with phosphate-buffered saline (PBS) for 5 min, followed by incubation in 10% sucrose/PBS for 3 h, and then in 25% sucrose/PBS overnight at 4°C. Afterward, the eyes were flash-frozen in a frozen section compound (Leica, Germany). Frozen tissue samples were sectioned at 12 *μ*m using a cryostat. For paraffin sections, enucleated eyes, lungs, and livers were fixed in 4% paraformaldehyde at 4°C for <24 h, and then embedded in paraffin. Paraffin-embedded tissue samples were sectioned at 3 *μ*m using a microtome. Tissue sections were stained with hematoxylin-eosin and analyzed using ImageJ software to measure ONL thickness. TUNEL staining was performed (TaKaRa Biochemicals, Japan) to detect apoptotic cells in the retina. For immunohistochemical analyses of PDE6β and rhodopsin, rabbit polyclonal PDE6β (ab5663; Abcam, UK) and 1D4 (ab3424; Abcam, UK) antibodies were used, respectively. Immunohistochemical analyses were performed using the VECTASTAIN Elite ABC kit (Vector Laboratories, UK).

### Quantitative Reverse transcriptase PCR (qPCR)

Total RNA was extracted from the retina or isolated photoreceptor cells using an RNeasy Mini Kit (Qiagen, Germany). Single-strand cDNA was synthesized using SuperScript III reverse transcriptase (Thermo Fisher Scientific, USA.). Using random primers (Thermo Fisher Scientific, USA.), qPCR was performed using the Fast SYBR Green Master Mix (Applied Biosystems, USA.) with the StepOnePlus real-time PCR system (Applied Biosystems, USA.). Relative mRNA expression levels were calculated using the comparative cycle threshold method and normalized to GAPDH mRNA levels. Primers for qPCR were as follows: Gene name: forward primer (5’ −;3’), reverse primer (5’−3’), and amplicon size (bp). *Gapdh*, TGACAATGAATACGGCTACAGCA, CTCCTGTTATTATGGGGGTCTGG (188) *Lpcat1*; TTCCCTGGGACCTCCTGATAAGG, GAGGAATGTGGAGCCAAGGTGAG, (186). *Rhodopsin*; CTCTTCATGGTCTTCGGAGGATT, GTGAAGACCACACCCATGATAGC, (230). *Fosb*; AGGCAGAGCTGGAGTCGGAGAT, GCCGAGGACTTGAACTTCACTTCG, (151). *Egr1*: CAACCCTATGAGCACCTGACCAC, GTCGTTTGGCTGGGATAACTCGT (95) *Egr2*; GCCAAGGCCGTAGACAAAATC, CCACTCCGTTCATCTGGTCA, (154). *Atf3*; TTTGCTAACCTGACACCCTTTG, AGAGGACATCCGATGGCAGA, (81). *Jun*; CCTTCTACGACGATGCCCTC, GGTTCAAGGTCATGCTCTGTTT, (102). *Junb*; TCACGACGACTCTTACGCAG, CCTTGAGACCCCGATAGGGA, (125). *Rpe65*; AGTTCCCCTGCAGTGATCGTTTC, ACCCCCATGCTTTCATTGGACTC, (178). *Chop*; CTGAGTCCCTGCCTTTCACCT, GCAGGGTCAAGAGTAGTGAAGGTTT, (166). *Scd*; ACTGGTTCCCTCCTGCAAG, GTGATCTCGGGCCCATTC, (199).

### Microsomal fraction preparation

The retina was homogenized in lysis buffer (100 mM Tris-HCl [pH 7.4], 300 mM sucrose) containing a 1X cOmplete protease inhibitor cocktail (Roche, Switzerland). After removing the tissue debris (800 × *g*, 10 min, 4°C) and microsomal fractions were collected by ultracentrifugation (100,000 × *g*, 60 min, 4°C). The pellets (microsomal fractions) were resuspended in ice-cold buffer containing 20 mM Tris-HCl (pH 7.4), 300 mM sucrose, and 1 mM EDTA. Protein concentrations were measured by the Bradford method using a protein assay (Bio-Rad, USA.).

### LC-MS/MS

For PC analyses, lipids were extracted from microsomal fractions or retinal OS using the Bligh & Dyer method (43). Subsequently, the extracted lipids were dried using a centrifugal evaporator and reconstituted in methanol. Lipid samples were then analyzed by LC-electrospray ionization-MS/MS with multiple reaction monitoring (MRM). LC-MS/MS analyses were performed using a Nexera UHPLC system and triple quadrupole mass spectrometer LCMS-8050 (Shimadzu Corp., Japan). The lipid samples (5 *μ*L/injection) were separated on an Acquity UPLC BEH C8 column (1.7 *μ*m, 2.1 × 100 mm, Waters) at a flow rate of 0.35 mL/min with a gradient of mobile phase A (5 mM NH4HCO3/water), B (acetonitrile), and C (isopropyl alcohol). The column oven temperature was set to 47°C. The gradient setting was as follows; time (%A/%B/%C): 0 min (50/45/5), 10 min (20/75/5), 20 min (20/50/30), 27.5 min (5/5/90), 28.5 min (5/5/90), 28.6 min (50/45/5) for Retina, and 0 min (75/20/5), 20 min (20/75/5), 40 min (20/5/75), 45 min (5/5/90), 50 min (5/5/90), 55 min (75/20/5) for the retinal OS. PC species were detected in positive ion mode following MRM transitions: (Q1, Q3), ([M + H]^+^, 184.1). The fatty acid composition of PC is denoted as Z XX:YY (where Z indicates the lipid class). XX and YY indicate the sum of the carbon numbers and double bonds, respectively.

For the analysis of ceramides, lipids were extracted from the mouse retina as previously described, with slight modifications (44). Briefly, retinas were homogenized in lysis buffer (100 mM Tris-HCl [pH 7.4], 300 mM sucrose) containing a 1X cOmplete protease inhibitor cocktail (Roche, Switzerland). After removing the tissue debris (800 × *g*, 10 min, 4°C) and 20 *μ*L of lysate was mixed with 30 *μ*L of PBS, 50 *μ*L of d18:1/12:0-ceramide (50 *μ*M in methanol), 750 *μ*L of *n*-butanol, 30 *μ*L of phosphate buffer (500 mM, pH 5.8), and 170 *μ*L of water. The samples were then vortexed for 5 min, sonicated for 3 min in a bath sonicator, and centrifuged at 1,000 × *g* for 5 min at 4°C. An upper phase (700 *μ*L) of the first butanol extract was transferred to a new tube. 350 *μ*L of ethyl acetate and 350 *μ*L of hexane were added to the remaining lower phase for the second extraction. After vortexing for 5 min at 4°C, 700 *μ*L of the upper phase (second extract) was combined with the first butanol extract and added to 700 *μ*L of methanol. The combined lipid extract (210 *μ*L,10%) was dried using a centrifugal evaporator, reconstituted in 60 *μ*L of solvent B (isopropyl alcohol with 0.2% formic acid and 0.028% ammonia), and then mixed with 90 *μ*L of solvent A (isopropyl alcohol/methanol/water (5:1:4) with 0.2% formic acid, 0.028% ammonia, and 5 *μ*M phosphoric acid). The samples were analyzed by LC-MS/MS with selected reaction monitoring (SRM). LC-MS/MS analysis was performed using an AQUITY UPLC system and a TSQ Vantage triple stage quadrupole mass spectrometer. The extracted lipids (5 *μ*L/injection) were separated on an Acclaim PepMap 100 C18 column (3 *μ*m, 1.0 × 150 mm, Thermo Fisher Scientific) at a flow rate of 0.045 mL/min with a linear gradient of solvent A and solvent B. The column temperature was 45°C. The gradient transition was as follows; time (%A/%B): 0 min (70/30), 2 min (50/50), 12 min (20/80), 12.5 min (5/95), 22.5 min (5/95), 23 min (95/5), 24 min (95/5), 26 min (70/30), 28 min (70/30). Ceramide species were detected in positive ion mode following MRM transitions (Q1, Q3), ([M – H_2_O + H]^+^, 264.3 or 266.3).

### Microarray

Total RNA from the whole retina or isolated photoreceptor cells was extracted using the RNeasy Mini Kit (QIAGEN, Germany). Total RNA was examined using the SurePrint G3 Mouse GE 8×60K Microarray (Agilent Technologies, USA.). Data were quantified using the Agilent Feature Extraction software (Agilent Technologies, USA.) and normalized by 75 percentile shift normalization using GeneSpring software (Agilent Technologies, USA.). The list of highly expressed genes in *Lpcat1* KO than in control with fold change > 1.5 or 2 and *P* < 0.05, was analyzed for common functions of altered genes using gene ontology (GO) terms using DAVID.

### Isolation of photoreceptor cells

Photoreceptor cells were isolated from the mouse retina as previously described, with slight modifications (17,45). Retinas were isolated from *Lpcat1* HZ and KO mice and dissociated in 500 *μ*L of 0.25% trypsin (Nacalai Tesque, Japan) in PBS for 15 min at 37°C. Then, 500 *μ*L of 20% fetal bovine serum (Thermo Fisher Scientific, USA.) and 1 *μ*L DNase I (Invitrogen, Germany) were added to the ice. Mechanical dissociation was performed by pipetting 20 times using a 1 mL tip. The cells were collected by centrifugation at 300 × *g* for 5 min. The retinal cell pellet was washed in 700 *μ*L of 2% bovine serum albumin (BSA)/ PBS and centrifuged at 300 × *g* for 5 min. The cell pellet was resuspended in 50 *μ*L of 2% BSA/PBS, and photoreceptor cells were labeled with PE rat anti-mouse CD73 antibody (TY/23, BD Pharmingen, USA.) for 30 min at 4°C. Cells were washed with 700 *μ*L of 2% BSA/PBS and centrifuged at 300 × *g* for 5 min. The cell pellet was resuspended in 80 *μ*L of 2% BSA/PBS and incubated with 20 *μ*L of anti-PE MicroBeads UltraPure (Miltenyi Biotec, USA.) for 15 min at 4°C. After washing with 700 *μ*L of 2% BSA/PBS and centrifugation at 300 × *g* for 5 min, the cell pellet was resuspended in 500 *μ*L of 2% BSA/PBS and filtered through a 35-*μ*m pre-separation filter (Corning, USA.). The CD73 positive cells were collected as photoreceptor cells using an autoMACS Pro Separator (Miltenyi Biotec, USA.).

### Retinal outer segment (OS) isolation

Mouse retinal OS was purified as described previously (46). Twelve mouse retinas were suspended in 120 *μ*L of 8% OptiPrep (Abbott Diagnostics Technologies AS, Norway) in Ringer’s buffer (130 mM NaCl, 3.6 mM KCl, 2.4 mM MgCl_2_, 1.2 mM CaCl_2_, 0.02 mM EDTA, 10 mM HEPES-NaOH, pH7.4). Retinal OS fractions were separated from the retina by vortexing for 1 min. The samples were then centrifuged at 200 × *g* for 1 min. The vortexing and sedimentation sequence was repeated six times to collect the remaining retinal OS fraction in the retinal pellets. The collected supernatants (retinal OS fractions) were combined (600 *μ*L) and overlaid onto a discontinuous gradient of OptiPrep™ (Veritas, Japan) (10%, 18%, 24%, and 30%) in Ringer’s buffer (1.8 mL each), and centrifuged at 25,900 × *g* for 30 min. After centrifugation, the retinal OS fraction (approximately two-thirds of the way from the top of the gradient) was harvested and diluted three times with Ringer’s buffer. The contaminated retinal fractions were removed by centrifugation at 500 × *g* for 3 min. The supernatant (pure retinal OS fraction) was precipitated by centrifugation at 26,500 × *g* for 30 min. The resultant retinal OS pellets (microsomal fractions) were resuspended in ice-cold buffer containing 20 mM Tris-HCl (pH 7.4), 300 mM sucrose, and 1 mM EDTA.

### Determination of cGMP levels

P10 mice were dark-adapted for 16 h, and retinas were dissected under dim red light and frozen in liquid nitrogen. Two frozen retinas were homogenized in 200 *μ*L of 5% trichloroacetic acid/water under dim red light and centrifuged at 1,500 × *g* for 10 min. The resultant supernatant was used for cGMP measurements. cGMP levels were determined using a cyclic GMP EIA kit (Cayman, USA.) according to the manufacturer’s instructions.

### Transmitted electron microscopy (TEM)

Enucleated eyes were immediately cut into hemispheres in a fixative containing 2% paraformaldehyde and 2.5% glutaraldehyde. The samples were then fixed in the same fixative for 2 h, followed by an additional fixation in the buffer containing 2% OsO4. The samples were dehydrated in a graded series of ethanol (70%-90%-100%) and embedded in an Epon 812 resin mixture (TAAB Laboratories Equipment Ltd., Berks, UK). Ultrathin sections were created using an ultramicrotome, stained with uranyl acetate and lead citrate, and observed under an electron microscope (Hitachi H-7500, Tokyo, Japan).

### Detection of mitochondrial reactive oxygen species (ROS)

ROS production was measured by flow cytometry using MitoSOX Red (Invitrogen, Germany). Retina was isolated from 3-week-old *Lpcat1* WT and KO mice and dissociated in 500 *μ*L of 0.25% trypsin (Nacalai Tesque, Japan) in PBS for 15 min at 37°C. Then, 500 *μ*L of 20% fetal bovine serum (Thermo Fisher Scientific, USA.) and 1 *μ*L DNase I (Invitrogen, Germany) were added to the ice. Mechanical dissociation was performed by pipetting 20 times using a 1 mL tip. The cells were collected by centrifugation at 300 × *g* for 5 min. The retinal cell pellet was washed in 700 *μ*L of 2% BSA/PBS and centrifuged at 300 × *g* for 5 min. The cell pellets were resuspended in 2% BSA/PBS. Photoreceptor cells were labeled with FITC anti-mouse CD73 antibody (TY/11.8, BioLegend, USA.). Dead cells were stained with LIVE/DEAD Fixable Far Red Dead Cell Stain kit (Invitrogen, Germany), and retinal cells were stained with 5 *μ*M MitoSOX Red for 15 min at 37°C. The cells were centrifuged at 300 × *g* for 5 min. The cell pellet was resuspended in 500 *μ*L of 2% BSA/PBS and filtered through a 35-*μ*m pre-separation filter (Corning, USA.). The fluorescence intensity was measured using a BD Accuri flow cytometer (BD Biosciences, USA.).

### Statistical Analysis

Unpaired t-tests were used when the two groups were compared. When two factors were present, a two-way ANOVA test was performed. Bonferroni’s post hoc test was used when the ANOVA showed significance. All statistical analyses were performed using GraphPad Prism9 (version 9.2.0).

## Data availability

The microarray data are deposited at GEO: GSE184815 (2-week-old retina).

The microarray data are deposited at GEO: GSE184817 (P8 photoreceptor cell).

## Acknowledgments

We are grateful to K. Yanagida (National Center for Global Health and Medicine) for their critical comments and valuable suggestions. We are grateful to T. Yokomizo (Juntendo University), K. Waku (Teikyo University), all members of our laboratories (National Center for Global Health and Medicine and University of Tokyo), and H. Koso (Institute of Medical Science, the University of Tokyo) for their valuable suggestions. We thank A. Tsuhako (Institute of Medical Science, University of Tokyo) for supporting retinal isolation. We thank M. Tamura-Nakano and C. Oyama (National Center for Global Health and Medicine) for supporting the histological analyses. We thank M. Matsumoto (National Center for Global Health and Medicine) for supporting the microarray analysis.

## Author Contributions

K.N. and D.H. designed the project, performed the experiments, interpreted the data, and wrote the manuscript. H.Sa. and M.S. performed the experiments and interpreted the data. S.W., T.S., and H.Sh. revised the manuscript. T.S. and H.Sh. supervised the project. All authors discussed the results and commented on the manuscript.

## Funding and additional information

This study was supported by AMED-CREST 21gm0910011(H.Sh.), AMED-P-CREATE 21cm0106116 (H.Sh.), AMED Program for Basic and Clinical Research on Hepatitis 21fk0210091h (H.Sh.) and Takeda Science Foundation 15668360 (T.S.).

## Abbreviations

The abbreviations used are

AGPAT3: 1-acylglycerol-3-phosphate *O*-acyltransferase 3
AMD: age-related macular degeneration
ANOVA: analysis of variance
AP-1: Activator protein-1
CHOP: CCAAT-enhancer-binding protein homologous protein
CNG: cyclic nucleotide-gated
Crb1: Crumbs homolog 1
DAVID: The Database for Annotation Visualization and Integrated Discovery
DEGs: differentially expressed genes
DHA: docosahexaenoic acid
DPPC: dipalmitoyl PC
ER: endoplasmic reticulum
FA: fatty acid
GCL: ganglion cell layer
GO: gene ontology
HZ: heterozygote
IEGs: immediate early genes
INL: inner nuclear layer
KO: knockout
LPAAT3: lysophosphatidic acid acyltransferase 3
LPC: lysophosphatidylcholine
LPCAT1: lysophosphatidylcholine acyltransferase 1
LPLATs: lysophospholipid acyltransferases
MRM: multiple reaction monitoring
ONL: outer nuclear layer
OS: outer segment
P8: postnatal day 8
PC: phosphatidylcholine
PDE6β: phosphodiesterase 6β
PUFA: polyunsaturated FA
rd: retinal degeneration
ROS: reactive oxygen species
RPE: retinal pigment epithelial
SCD: stearoyl-CoA desaturase
SFA: saturated FA
SM: sphingomyelin
SRM: selected reaction monitoring
TEM: transmitted electron microscopy
TUNEL: terminal deoxynucleotidyl transferase dUTP nick-end labeling.

## Supporting information

### Supplementary methods

#### Immunoblot

Protein samples were resolved on 4% / 10% SDS-polyacrylamide gels and transferred to nitrocellulose membranes (GE Healthcare, USA.) using a Trans-Blot Turbo (Bio-Rad, USA). Membranes were blocked with blocking buffer (5% skim milk in Tris-based buffer with 0.1% polyoxyethylene(20) sorbitan monolaurate (Wako, Japan)) at room temperature for 1 h. The membrane was incubated with rabbit-anti-LPCAT1 antibody (47) or mouse-anti-β-actin (GE Healthcare) at 4°C for 16 hours. The membrane was washed three times with wash buffer (Tris-based buffer with 0.1% polyoxyethylene(20) sorbitan monolaurate) for 5 min, and incubated with anti-rabbit IgG antibody conjugated to horseradish (GE Healthcare, USA.) or anti-mouse IgG antibody conjugated to horseradish (GE Healthcare) at room temperature for 1 h. The membrane was washed three times with wash buffer for 5 min each and developed using ECL reagent (GE Healthcare, USA.). Immunoreactive proteins were visualized using ImageQuant LAS500 (GE Healthcare, USA.).

### Measurement of total sphingomyelin (SM) levels

Retinal lipids were separated using thin layer chromatography (TLC) to determine the total SM levels. Lipids were extracted from the frozen retina as follows: The frozen retinas were sonicated in 200 *μ*L of methanol. Then, the retinal lipids in the methanol suspension were further extracted using the Bligh and Dyer method (43). The extracted lipids were treated with KOH (final 0.125N), followed by 15 min of heating at 60°C. Lipid solutions were spotted (1 *μ*L) onto chromatoplates precoated with silica gel 60 (Merck, Germany). The developing solvent system used in the TLC experiment was chloroform: methanol: acetic acid: water (25:20:4:2, v/v/v/v). Lipid spots were detected with primulin and then visualized using ImageQuant LAS500 (GE Healthcare, USA.). SM was identified by comparison with the spot of authentic SM (860062; Avanti Polar Lipids, Inc., USA.). The signal intensity of SM was quantified using Image J software.

---

**Figure S1.**
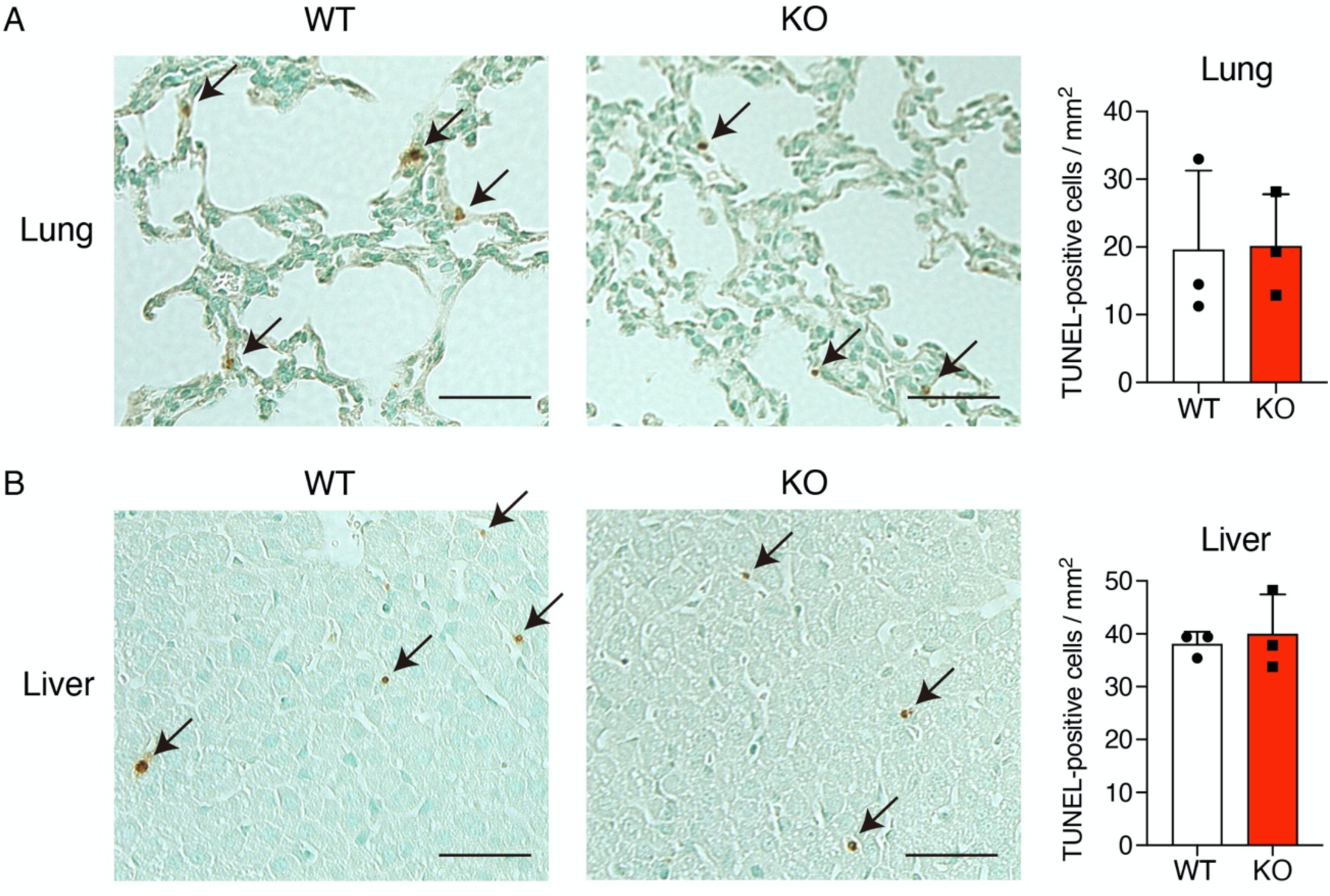
Apoptotic cells of *Lpcat1* wild-type (WT) and knockout (KO) lung and liver. Representative images of terminal deoxynucleotidyl transferase dUTP nick-end labeling (TUNEL) staining of 2-week-old *Lpcat1* WT and KO lung (A) and liver (B). Nuclei were stained with Methyl Green. Arrows indicated TUNEL-positive nuclei, which is detected by 3,3’-diaminobenzidine (DAB, shown in brown). Right bar graphs showed the average number of TUNEL-positive cells per 1 mm^2^ in randomly selected areas. Data are shown as mean + SD of independent experiments (n=3 for each group). Scale bars are 50 *μ*m.

**Figure S2.**
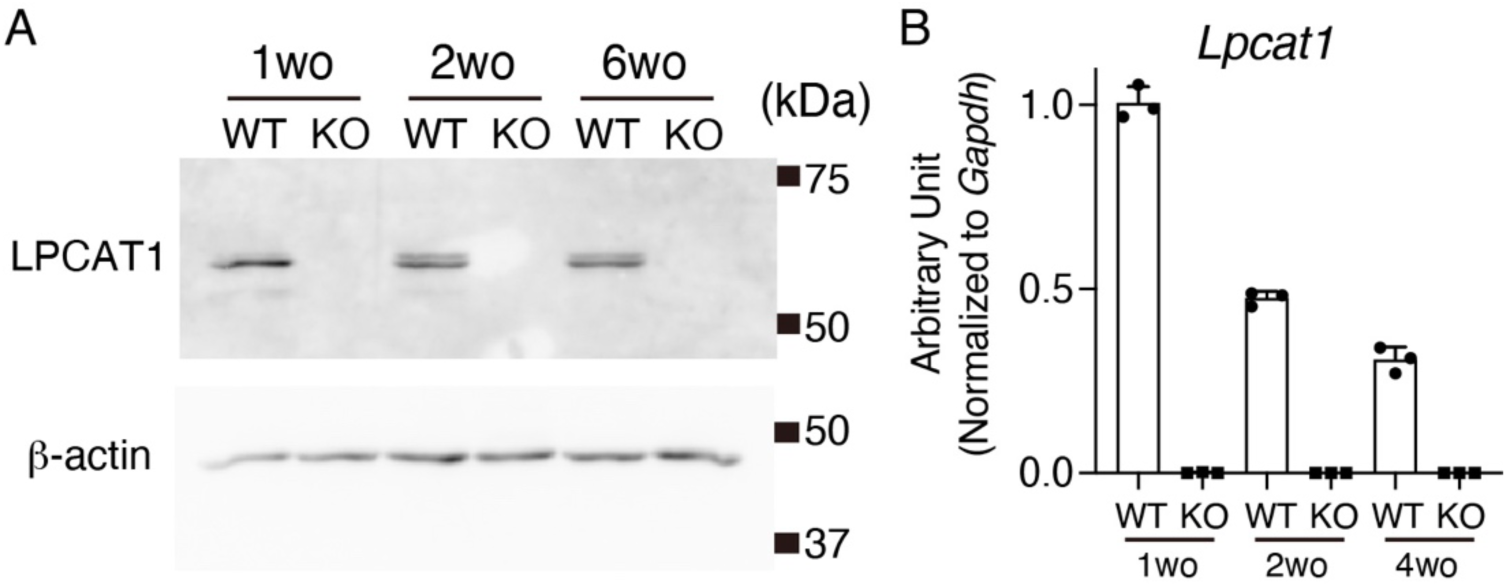
The expression of LPCAT1 mRNA and protein during retinal maturation in *Lpcat1* wild-type (WT) and knockout (KO) retina. A, Representative image of immunoblot analysis of the amount of LPCAT1 in 1-, 2- and 6-week-old (wo) *Lpcat1* wild-type (WT) and knockout (KO) retina. Similar results were obtained in three independent experiments. β-actin was used as a loading control. B, LPCAT1 mRNA expression in 1-, 2- and 4-week-old *Lpcat1* WT and KO retina. The expression level was normalized by the glyceraldehyde 3-phosphate dehydrogenase (*Gapdh*) gene. Data are shown as mean + SD of independent experiments (n=3 for each group).

**Figure S3.**
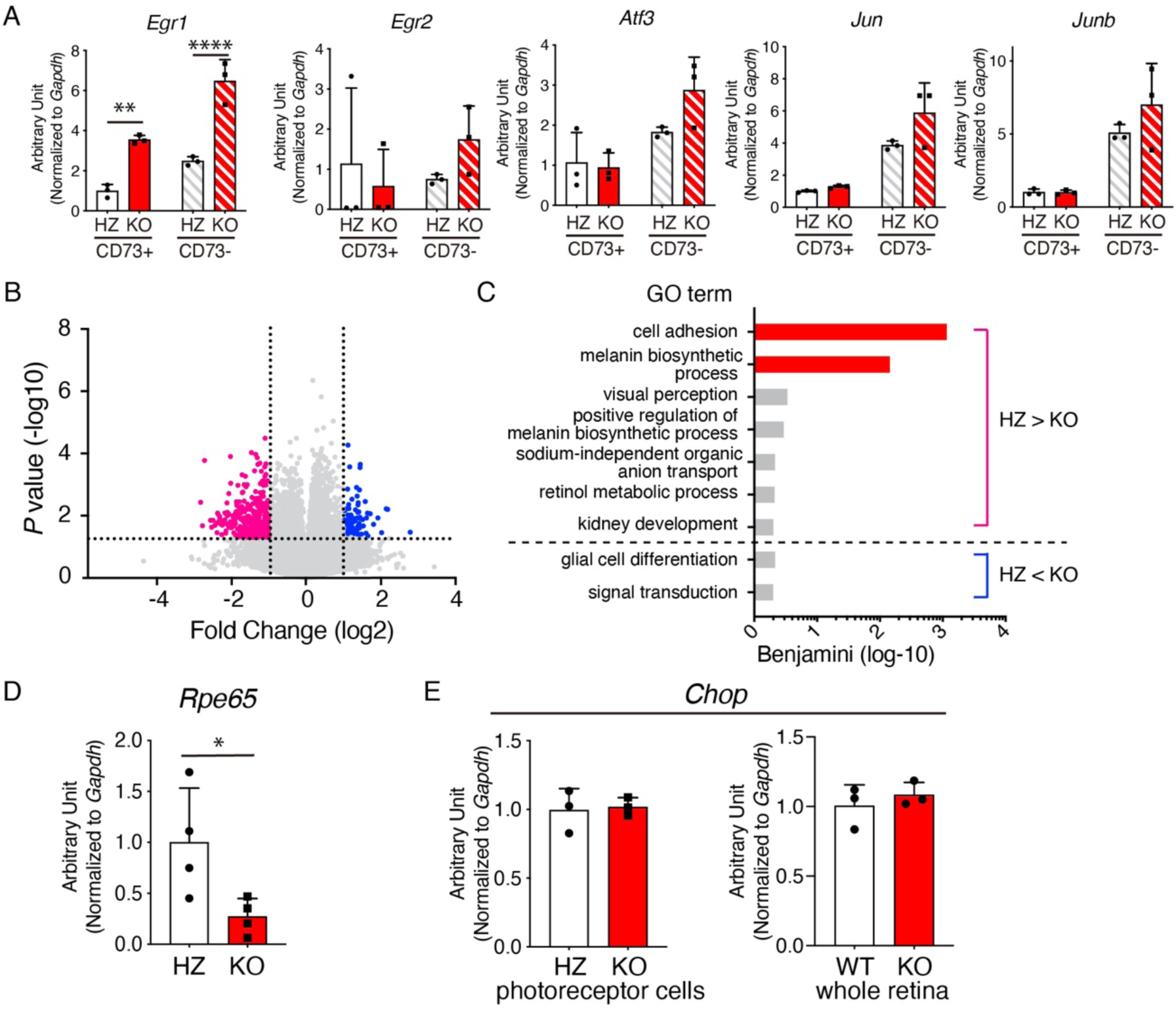
Gene expression analyses of *Lpcat1* knockout (KO) retinal cells. A, Gene expression of immediate early genes in postnatal day 14 (P14) *Lpcat1* heterozygote (HZ) and KO of CD73 positive (photoreceptor cells) and negative retinal cells. Data are shown as mean + SD of independent experiments (n=3 for each group). Significance is based on two-way ANOVA followed by Bonferroni’s multiple comparisons test (\*\**P* < 0.01, \*\*\*\**P* < 0.0001). B and C, Transcriptomic analysis of P8 *Lpcat1* HZ and KO photoreceptor cells. B, Volcano plot of differentially expressed genes (DEGs) between P8 *Lpcat1* HZ and KO photoreceptor cells. DEGs (Fold change > 2 or < -2, *P* < 0.05) are highlighted in blue; increased in *Lpcat1* KO, and magenta; decreased in *Lpcat1* KO (n=4 for each group). C, Functional annotation of down- and upregulated genes in *Lpcat1* KO photoreceptor cells. The top gene ontology (GO) terms from “biological processes” are shown (Benjamini-corrected *P* < 0.5). D, RPE65 mRNA expression in P8 *Lpcat1* HZ and KO photoreceptor cells. Data are shown as mean + SD of independent experiments (n=4 for each group). Significance is based on unpaired *t*-test (\**P* < 0.05). E, CHOP mRNA expression in P14 *Lpcat1* HZ and KO photoreceptor cells and P14 *Lpcat1* wild-type (WT) and KO whole retina. Data are shown as mean + SD of independent experiments (n=3 for each group). A, D, and E, The expression level was normalized by glyceraldehyde 3-phosphate dehydrogenase (*Gapdh*).

**Figure S4.**
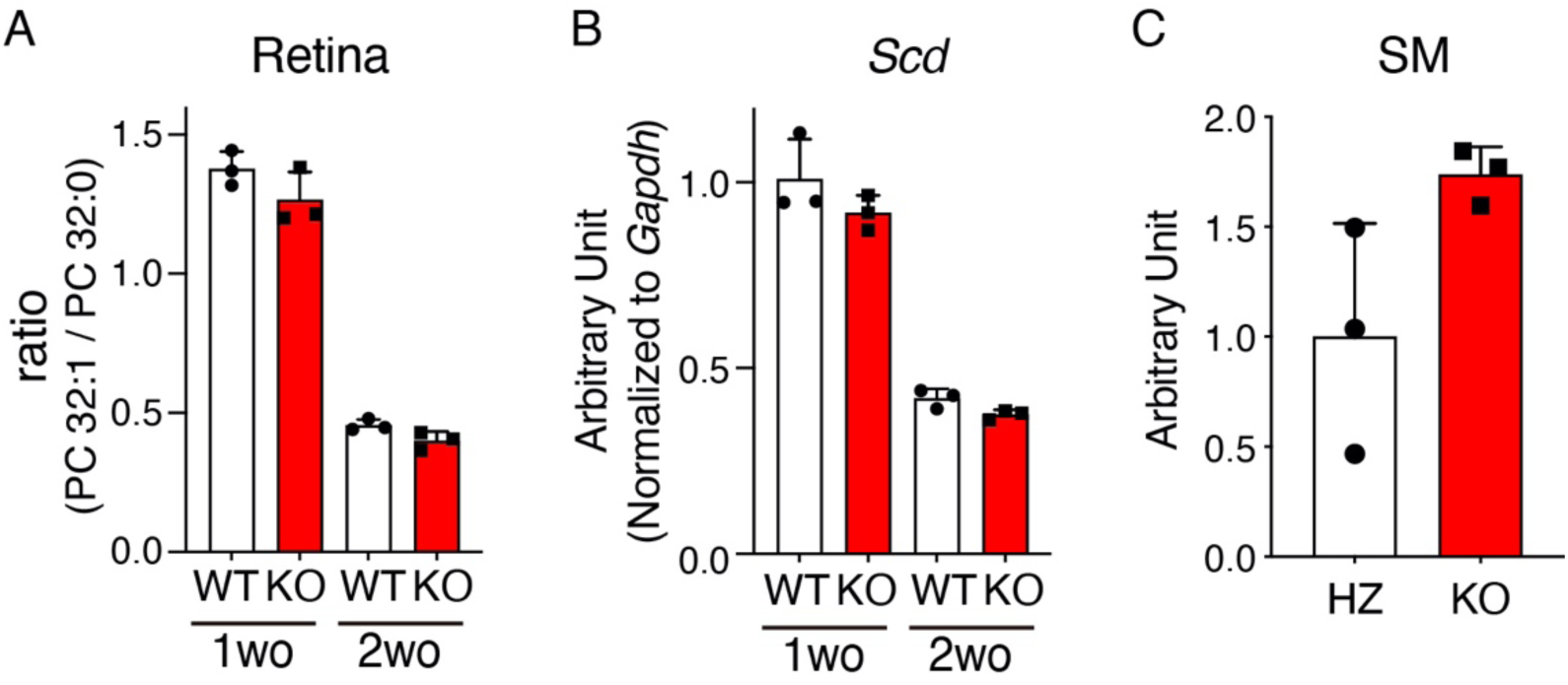
Fatty acid-related factors during retinal maturation. A, Relative ratio of area values of PC 32:1 to PC 32:0 in 1- and 2-week-old (wo) *Lpcat1* wild-type (WT) and knockout (KO) retina. Data are shown as mean + SD of independent experiments (n=3 for each group). B, Gene expression of *Scd* at mRNA level in 1- and 2-week-old *Lpcat1* WT and KO retina. The expression level was normalized by glyceraldehyde 3-phosphate dehydrogenase (*Gapdh*). Data are shown as mean + SD of independent experiments (n=3 for each group). C, Total sphingomyelin (SM) levels in 2-week-old *Lpcat1* heterozygote (HZ) and KO retina. Lipids extracted retinal tissues were separated with thin layer chromatography (TLC) and stained with primulin. The spot of SM was identified using authentic SM. The signal intensity of SM was quantified by Image J software. Data are shown as mean + SD of independent experiments (n=3 for each group).

**Figure S5.**
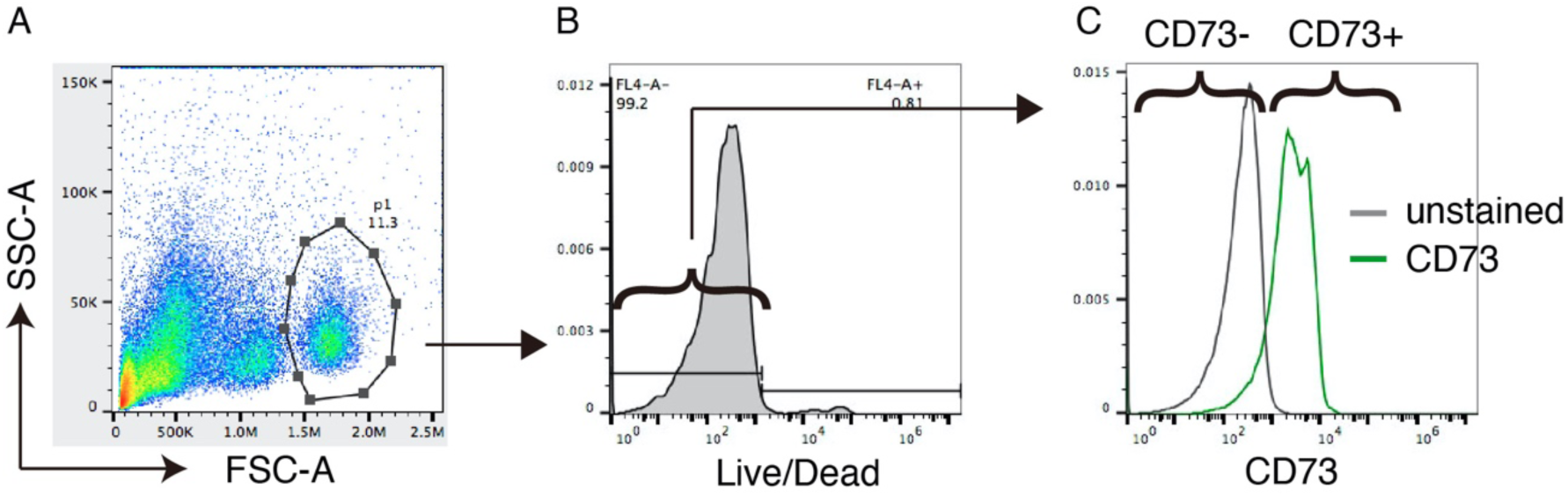
Gating strategy of retinal cells. A-C, Retinal cells were labeled with Live/Dead fluorescent dye and anti-CD73 antibody. A, Retinal cells (inside the indicated gate) were separated from cellular debris by forward versus side scatter (FSC vs SSC) gating. B, Living retinal cells were selected as Live/Dead staining-negative cells. C, Retinal cells were further separated into CD73-positive (photoreceptor) and -negative cells.

**Table S1.**
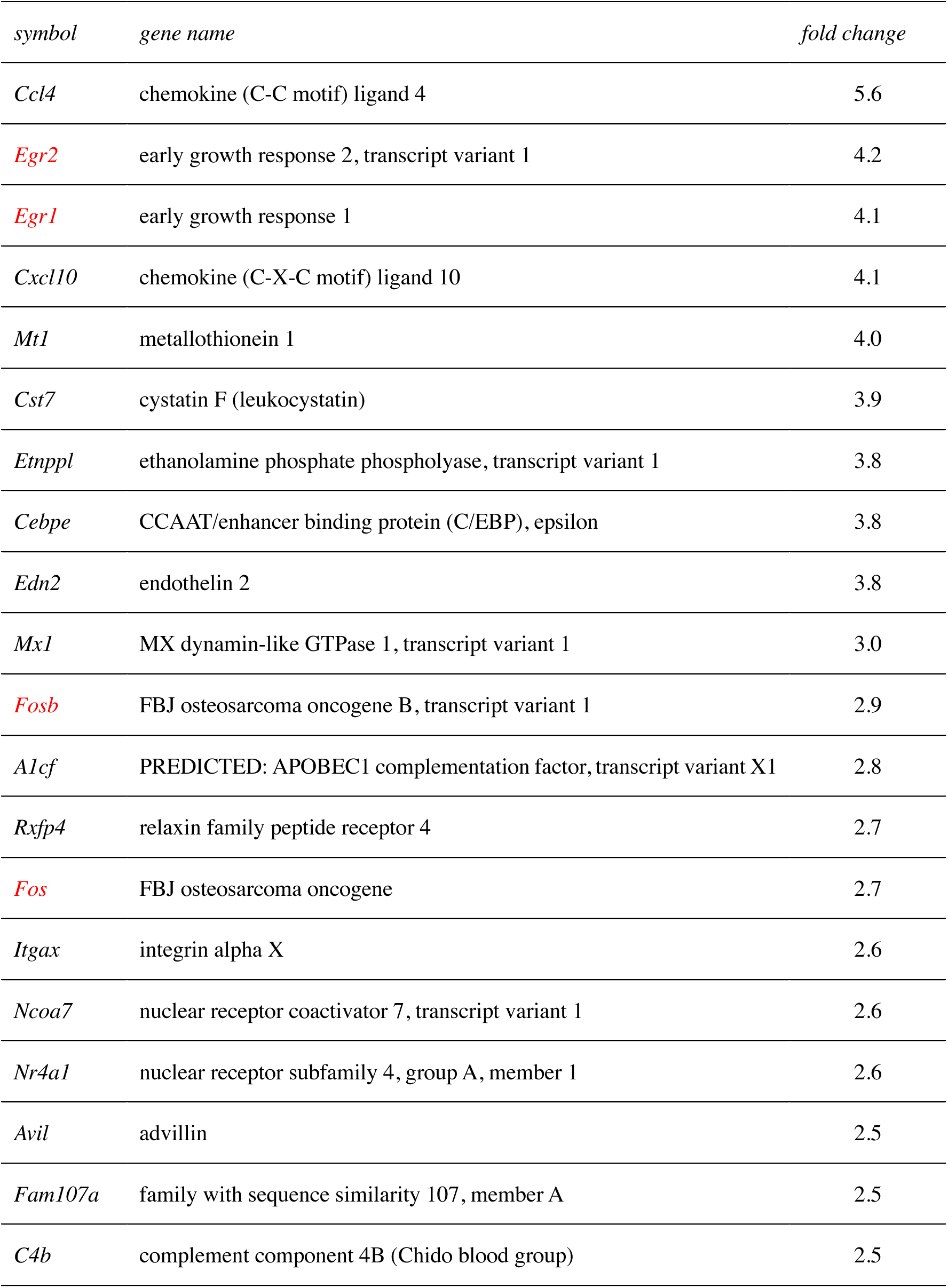
List of upregulated genes in 2-week-old *Lpcat1* knockout retina. Immediate early genes (IEGs) are shown in red.

## References

1. Kennedy, E. P., and Weiss, S. B. (1956) The function of cytidine coenzymes in the biosynthesis of phospholipides. J Biol Chem 222, 193–214

2. Lands, W. E. (1958) Metabolism of glycerolipides; a comparison of lecithin and triglyceride synthesis. J Biol Chem 231, 883–888

3. Shimizu, T. (2009) Lipid mediators in health and disease: enzymes and receptors as therapeutic targets for the regulation of immunity and inflammation. Annu Rev Pharmacol Toxicol 49, 123–150

4. Nakanishi, H., Shindou, H., Hishikawa, D., Harayama, T., Ogasawara, R., Suwabe, A., Taguchi, R., and Shimizu, T. (2006) Cloning and characterization of mouse lung-type acyl-CoA:lysophosphatidylcholine acyltransferase 1 (LPCAT1). Expression in alveolar type II cells and possible involvement in surfactant production. J Biol Chem 281, 20140–20147

5. Shindou, H., and Shimizu, T. (2009) Acyl-CoA:lysophospholipid acyltransferases. J Biol Chem 284, 1–5

6. Harayama, T., Eto, M., Shindou, H., Kita, Y., Otsubo, E., Hishikawa, D., Ishii, S., Sakimura, K., Mishina, M., and Shimizu, T. (2014) Lysophospholipid acyltransferases mediate phosphatidylcholine diversification to achieve the physical properties required in vivo. Cell Metab 20, 295–305

7. Bridges, J. P., Ikegami, M., Brilli, L. L., Chen, X., Mason, R. J., and Shannon, J. M. (2010) LPCAT1 regulates surfactant phospholipid synthesis and is required for transitioning to air breathing in mice. J Clin Invest 120, 1736–1748

8. Bi, J., Ichu, T. A., Zanca, C., Yang, H., Zhang, W., Gu, Y., Chowdhry, S., Reed, A., Ikegami, S., Turner, K. M., Zhang, W., Villa, G. R., Wu, S., Quehenberger, O., Yong, W. H., Kornblum, H. I., Rich, J. N., Cloughesy, T. F., Cavenee, W. K., Furnari, F. B., Cravatt, B. F., and Mischel, P. S. (2019) Oncogene Amplification in Growth Factor Signaling Pathways Renders Cancers Dependent on Membrane Lipid Remodeling. Cell Metab 30, 525–538 e528

9. Friedman, J. S., Chang, B., Krauth, D. S., Lopez, I., Waseem, N. H., Hurd, R. E., Feathers, K. L., Branham, K. E., Shaw, M., Thomas, G. E., Brooks, M. J., Liu, C., Bakeri, H. A., Campos, M. M., Maubaret, C., Webster, A. R., Rodriguez, I. R., Thompson, D. A., Bhattacharya, S. S., Koenekoop, R. K., Heckenlively, J. R., and Swaroop, A. (2010) Loss of lysophosphatidylcholine acyltransferase 1 leads to photoreceptor degeneration in rd11 mice. Proc Natl Acad Sci U S A 107, 15523–15528

10. Dai, X., Han, J., Qi, Y., Zhang, H., Xiang, L., Lv, J., Li, J., Deng, W. T., Chang, B., Hauswirth, W. W., and Pang, J. J. (2014) AAV-mediated lysophosphatidylcholine acyltransferase 1 (Lpcat1) gene replacement therapy rescues retinal degeneration in rd11 mice. Invest Ophthalmol Vis Sci 55, 1724–1734

11. Hamano, F., Kuribayashi, H., Iwagawa, T., Tsuhako, A., Nagata, K., Sagara, H., Shimizu, T., Shindou, H., and Watanabe, S. (2021) Mapping membrane lipids in the developing and adult mouse retina under physiological and pathological conditions using mass spectrometry. J Biol Chem 296, 100303

12. Shindou, H., Koso, H., Sasaki, J., Nakanishi, H., Sagara, H., Nakagawa, K. M., Takahashi, Y., Hishikawa, D., Iizuka-Hishikawa, Y., Tokumasu, F., Noguchi, H., Watanabe, S., Sasaki, T., and Shimizu, T. (2017) Docosahexaenoic acid preserves visual function by maintaining correct disc morphology in retinal photoreceptor cells. J Biol Chem 292, 12054–12064

13. Hamel, C. (2006) Retinitis pigmentosa. Orphanet J Rare Dis 1, 40

14. Lem, J., Krasnoperova, N. V., Calvert, P. D., Kosaras, B., Cameron, D. A., Nicolo, M., Makino, C. L., and Sidman, R. L. (1999) Morphological, physiological, and biochemical changes in rhodopsin knockout mice. Proc Natl Acad Sci U S A 96, 736–741

15. Fu, Z., Kern, T. S., Hellstrom, A., and Smith, L. E. H. (2021) Fatty acid oxidation and photoreceptor metabolic needs. J Lipid Res 62, 100035

16. Sheng, M., and Greenberg, M. E. (1990) The regulation and function of c-fos and other immediate early genes in the nervous system. Neuron 4, 477–485

17. Koso, H., Minami, C., Tabata, Y., Inoue, M., Sasaki, E., Satoh, S., and Watanabe, S. (2009) CD73, a novel cell surface antigen that characterizes retinal photoreceptor precursor cells. Invest Ophthalmol Vis Sci 50, 5411–5418

18. Akagi, S., Kono, N., Ariyama, H., Shindou, H., Shimizu, T., and Arai, H. (2016) Lysophosphatidylcholine acyltransferase 1 protects against cytotoxicity induced by polyunsaturated fatty acids. The FASEB Journal 30, 2027–2039

19. Mendes, H. F., van der Spuy, J., Chapple, J. P., and Cheetham, M. E. (2005) Mechanisms of cell death in rhodopsin retinitis pigmentosa: implications for therapy. Trends Mol Med 11, 177–185

20. Chang, B., Hawes, N. L., Pardue, M. T., German, A. M., Hurd, R. E., Davisson, M. T., Nusinowitz, S., Rengarajan, K., Boyd, A. P., Sidney, S. S., Phillips, M. J., Stewart, R. E., Chaudhury, R., Nickerson, J. M., Heckenlively, J. R., and Boatright, J. H. (2007) Two mouse retinal degenerations caused by missense mutations in the beta-subunit of rod cGMP phosphodiesterase gene. Vision Res 47, 624–633

21. Arshavsky, V. Y., Lamb, T. D., and Pugh, E. N., Jr. (2002) G proteins and phototransduction. Annu Rev Physiol 64, 153–187

22. Lolley, R. N., and Farber, D. B. (1976) Abnormal guanosine 3’, 5’-monophosphate during photoreceptor degeneration in the inherited retinal disorder of C3H/HeJ mice. Ann Ophthalmol 8, 469–473

23. Chang, B., Hawes, N. L., Hurd, R. E., Davisson, M. T., Nusinowitz, S., and Heckenlively, J.R. (2002) Retinal degeneration mutants in the mouse. Vision Research 42, 517–525

24. Estadella, D., da Penha Oller do Nascimento, C.M., Oyama, L. M., Ribeiro, E. B., Damaso, A.R., and de Piano, A. (2013) Lipotoxicity: effects of dietary saturated and transfatty acids. Mediators Inflamm 2013, 137579

25. Scaglia, N., and Igal, R. A. (2005) Stearoyl-CoA desaturase is involved in the control of proliferation, anchorage-independent growth, and survival in human transformed cells. J Biol Chem 280, 25339–25349

26. Chen, L., Ren, J., Yang, L., Li, Y., Fu, J., Li, Y., Tian, Y., Qiu, F., Liu, Z., and Qiu, Y. (2016) Stearoyl-CoA desaturase-1 mediated cell apoptosis in colorectal cancer by promoting ceramide synthesis. Sci Rep 6, 19665

27. Egnatchik, R. A., Leamy, A. K., Noguchi, Y., Shiota, M., and Young, J. D. (2014) Palmitate-induced activation of mitochondrial metabolism promotes oxidative stress and apoptosis in H4IIEC3 rat hepatocytes. Metabolism 63, 283–295

28. Ariyama, H., Kono, N., Matsuda, S., Inoue, T., and Arai, H. (2010) Decrease in membrane phospholipid unsaturation induces unfolded protein response. J Biol Chem 285, 22027–22035

29. Volmer, R., and Ron, D. (2015) Lipid-dependent regulation of the unfolded protein response. Curr Opin Cell Biol 33, 67–73

30. Harayama, T., and Riezman, H. (2018) Understanding the diversity of membrane lipid composition. Nat Rev Mol Cell Biol 19, 281–296

31. Uematsu, M., and Shimizu, T. (2021) Raman microscopy-based quantification of the physical properties of intracellular lipids. Communi Biol 4, (in press)

32. Sander, C. L., Sears, A. E., Pinto, A. F. M., Choi, E. H., Kahremany, S., Gao, F., Salom, D., Jin, H., Pardon, E., Suh, S., Dong, Z., Steyaert, J., Saghatelian, A., Skowronska-Krawczyk, D., Kiser, P. D., and Palczewski, K. (2021) Nano-scale resolution of native retinal rod disk membranes reveals differences in lipid composition. J Cell Biol 220

33. Joyal, J. S., Sun, Y., Gantner, M. L., Shao, Z., Evans, L. P., Saba, N., Fredrick, T., Burnim, S., Kim, J. S., Patel, G., Juan, A. M., Hurst, C. G., Hatton, C. J., Cui, Z., Pierce, K. A., Bherer, P., Aguilar, E., Powner, M. B., Vevis, K., Boisvert, M., Fu, Z., Levy, E., Fruttiger, M., Packard, A., Rezende, F. A., Maranda, B., Sapieha, P., Chen, J., Friedlander, M., Clish, C. B., and Smith, L.E. (2016) Retinal lipid and glucose metabolism dictates angiogenesis through the lipid sensor Ffar1. Nat Med 22, 439–445

34. Pan, W. W., Wubben, T. J., and Besirli, C. G. (2021) Photoreceptor metabolic reprogramming: current understanding and therapeutic implications. Commun Biol 4, 245

35. Flowers, M. T., and Ntambi, J. M. (2008) Role of stearoyl-coenzyme A desaturase in regulating lipid metabolism. Curr Opin Lipidol 19, 248–256

36. Cheng, X., Geng, F., Pan, M., Wu, X., Zhong, Y., Wang, C., Tian, Z., Cheng, C., Zhang, R., Puduvalli, V., Horbinski, C., Mo, X., Han, X., Chakravarti, A., and Guo, D. (2020) Targeting DGAT1 Ameliorates Glioblastoma by Increasing Fat Catabolism and Oxidative Stress. Cell Metab 32, 229–242 e228

37. Listenberger, L. L., Han, X., Lewis, S. E., Cases, S., Farese, R. V., Jr., Ory, D. S., and Schaffer, J. E. (2003) Triglyceride accumulation protects against fatty acid-induced lipotoxicity. Proc Natl Acad Sci U S A 100, 3077–3082

38. Geller, S. F., and Stone, J. (2003) Quantitative PCR analysis of FosB mRNA expression after short duration oxygen and light stress. Adv Exp Med Biol 533, 249–257

39. Shaulian, E., and Karin, M. (2001) AP-1 in cell proliferation and survival. Oncogene 20, 2390–2400

40. Yanagida, K., Engelbrecht, E., Niaudet, C., Jung, B., Gaengel, K., Holton, K., Swendeman, S., Liu, C. H., Levesque, M. V., Kuo, A., Fu, Z., Smith, L. E. H., Betsholtz, C., and Hla, T. (2020) Sphingosine 1-Phosphate Receptor Signaling Establishes AP-1 Gradients to Allow for Retinal Endothelial Cell Specialization. Dev Cell 52, 779–793 e777

41. Grunin, M., Hagbi-Levi, S., Rinsky, B., Smith, Y., and Chowers, I. (2016) Transcriptome Analysis on Monocytes from Patients with Neovascular Age-Related Macular Degeneration. Sci Rep 6, 29046

42. Mattapallil, M. J., Wawrousek, E. F., Chan, C. C., Zhao, H., Roychoudhury, J., Ferguson, T. A., and Caspi, R. R. (2012) The Rd8 mutation of the Crb1 gene is present in vendor lines of C57BL/6N mice and embryonic stem cells, and confounds ocular induced mutant phenotypes. Invest Ophthalmol Vis Sci 53, 2921–2927

43. Bligh, E. G., and Dyer, W. J. (1959) A rapid method of total lipid extraction and purification. Can J Biochem Physiol 37, 911–917

44. Ogiso, H., Taniguchi, M., Araya, S., Aoki, S., Wardhani, L. O., Yamashita, Y., Ueda, Y., and Okazaki, T. (2014) Comparative Analysis of Biological Sphingolipids with Glycerophospholipids and Diacylglycerol by LC-MS/MS. Metabolites 4, 98–114

45. Eberle, D., Schubert, S., Postel, K., Corbeil, D., and Ader, M. (2011) Increased integration of transplanted CD73-positive photoreceptor precursors into adult mouse retina. Invest Ophthalmol Vis Sci 52, 6462–6471

46. Liang, Y., Fotiadis, D., Filipek, S., Saperstein, D. A., Palczewski, K., and Engel, A. (2003) Organization of the G protein-coupled receptors rhodopsin and opsin in native membranes. J Biol Chem 278, 21655–21662

47. Harayama, T., Shindou, H., Ogasawara, R., Suwabe, A., and Shimizu, T. (2008) Identification of a novel noninflammatory biosynthetic pathway of platelet-activating factor. J Biol Chem 283, 11097–11106

